# Functional divergence of two duplicated *Fertilization Independent Endosperm* genes in rice with respect to seed development

**DOI:** 10.1101/714444

**Authors:** Meiyao Pan, Xiaojun Cheng, E Zhiguo, Baixiao Niu, Chen Chen

## Abstract

Fertilization Independent Endosperm (FIE) is an essential member of Polycomb Repression Complex 2 (PRC2) that plays important roles in the developmental regulation of plants. *OsFIE1* and *OsFIE2* are two *FIE* homologs in the rice genome. Here, we showed that *OsFIE1* probably duplicated from *OsFIE2* after the origin of the tribe Oryzeae, but has a specific expression pattern and methylation landscape. During evolution, *OsFIE1* underwent a less intensive purifying selection than did *OsFIE2*. The mutant *osfie1* produced smaller seeds and displayed reduced dormancy, indicating that *OsFIE1* predominantly functions in late seed development. Ectopic expression of *OsFIE1*, but not *OsFIE2*, was deleterious to vegetative growth in a dosage-dependent manner. The newly evolved N-terminal tail of OsFIE1 was probably not the cause of the adverse effects on vegetative growth. The CRISPR/Cas9-derived mutant *osfie2* exhibited impaired cellularization of the endosperm, which suggested that *OsFIE2* is indispensable for early seed development as a positive regulator of cellularization. Autonomous endosperm was observed in both *OsFIE2*^+−^ and *osfie1*/*OsFIE2*^+−^ but at a very low frequency. Although OsFIE1-PRC2 exhibited H3K27me3 methyltransferase ability in plants, OsFIE1-PRC2 is likely to be less important for development in rice than is OsFIE2-PRC2. Our findings revealed the functional divergence of *OsFIE1* and *OsFIE2* and shed light on their distinct evolution following duplication.

## Introduction

The endosperm is a product of double fertilization in higher plants (Olsen, 2001). It is a triploid tissue, produced by fusion between a central cell (2n) and a sperm cell (n). The primary endosperm cell usually undergoes several rounds of mitotic division uncoupled from cytokinesis, which results in a multinucleate cell with many free nuclei, termed a syncytium (Olsen, 2001; Wu *et al.*, 2016). Then, the multiple nuclei start to cellularize and generate the first layer of endosperm cells (Olsen, 2001; Wu *et al.*, 2016). Subsequent cell divisions allow the endosperm cells to fill the seed. The fate of the endosperm is different in dicots and monocots; the endosperm is usually consumed in dicots during seed development, while it is retained in mature seeds of monocots. However, the developmental process of endosperm is quite highly conserved in plants (Olsen, 2004; Agarwal *et al.*, 2011). As a nutritional supply tissue, the endosperm is indispensable for embryo development. Either delayed or accelerated cellularization can lead to seed failure in interploidy and interspecific hybrids (Walia *et al.*, 2009; Ishikawa *et al.*, 2011; Kradolfer *et al.*, 2013; Tonosaki *et al.*, 2018). Developmental defects in the endosperm will cause embryo abortion, which has been found to occur in many plant species, where it acts as an important reproductive barrier (Chen *et al.*, 2016b).

Polycomb repression complex 2 (PRC2) plays an important role in developmental regulation by controlling epigenetic modification of the genome (Mozgova & Hennig, 2015; Mozgova *et al.*, 2015). PRC2 has methyltransferase activity for the methylation of Lys27 in histone H3 (H3K27) (Cao *et al.*, 2002; Nallamilli *et al.*, 2013). The major components of PRC2 are highly conserved in animals and plants (Tonosaki & Kinoshita, 2015), both of which include four group members: WD40 protein p55 (p55), Suppressor of Zeste 12 [Su(z)12], Enhancer of Zeste [E(z)], and extra sex combs (ESC). Fertilization Independent Seed (FIS)-PRC2 of Arabidopsis, composed of FIS2 (a Su(z)12 member), Fertilization Independent Endosperm (FIE, a (ESC) member), MEDEA (MEA, a E(z) member) and Multicopy Suppressors of IRA 1 (MSI1, a p55 member), acts as a key regulator of seed development (Mozgova *et al.*, 2015). Mutation of these genes results in autonomous endosperm, developing without fertilization (Ohad *et al.*, 1996; Chaudhury *et al.*, 1997; Kiyosue *et al.*, 1999). In addition, FIS-PRC2 is indispensable for the transition from the syncytium to cellularized cells in Arabidopsis (Grossniklaus *et al.*, 1998; Luo *et al.*, 2000; Vinkenoog *et al.*, 2000; Hennig, 2003).

Different plant species have evolved distinct members of each PRC2 group (Tonosaki & Kinoshita, 2015). For example, rice lacks MEA that belongs to the E(z) group and lacks FIS2 that belongs to the Su(z)12 group (Luo *et al.*, 2009). FIE is the only ESC member that encoded by the Arabidopsis genome. However, species of the Poaceae usually have multiple FIE homologs (Danilevskaya *et al.*, 2003a; Luo *et al.*, 2009; Kapazoglou *et al.*, 2010). OsFIE1 and OsFIE2, which show marked similarities, are two FIE homologs encoded by the rice genome (Luo *et al.*, 2009). To date, the mechanism by which PRC2 regulates endosperm development in species other than Arabidopsis is largely unknown. Overexpression of *OsFIE1* led to a phenotype of short stature, low seed-setting rate and small seeds (Zhang *et al.*, 2012; Folsom *et al.*, 2014), but no altered phenotype or a very mild phenotype change was exhibited by the RNA interference (RNAi)-mediated knock-down plant or T-DNA mutant of *OsFIE1* according to previous reports (Yang *et al.*, 2013; Huang *et al.*, 2016). On the other hand, *OsFIE2* RNAi plants displayed phenotypes similar to those of the *OsFIE1*-overexpressors, such as dwarfism and reduced seed-setting rate (Nallamilli *et al.*, 2013; Li *et al.*, 2014; Liu *et al.*, 2016). Some inconsistent findings have been reported according to previous studies. For example, Li *et al.* (2014) found that knock-down of *OsFIE2* led to an autonomous endosperm but Nallamilli *et al.* (2013) did not observe such an effect. Luo *et al.* (2009) found that a T-DNA mutant line of *OsFIE1* did not cause any morphological changes, whereas Huang *et al.* (2016) reported that RNAi of *OsFIE1* reduced seed size and delayed embryo development.

*MEA* and *FIS2* are recently evolved PRC2 members as a result of gene duplication in *Arabidopsis* (Spillane *et al.*, 2007). Evolutionary analysis revealed that *MEA* had undergone natural selection in the promoter region (Kawabe *et al.*, 2007; Spillane *et al.*, 2007; Miyake *et al.*, 2009). Currently, our understanding of how the PRC2 members evolved and how they contribute to adaptation in plants other than Arabidopsis is limited (Furihata *et al.*, 2016). Given the strong similarities between *OsFIE1* and *OsFIE2*, we believed that it would not be a good strategy to use RNAi lines for functional analysis, owing to the spatiotemporal overlaps of expression of the two genes in seeds (Li *et al.*, 2014). Therefore, systemic analysis of *OsFIEs* using null mutants is preferred to achieve a better understanding of the PRC2-regulated endosperm development and to clarify the controversies that has been reported. In the present study, we combined the use of evolutionary and genetic approaches to dissect the functional divergence of *OsFIE1* and *OsFIE2*. Our results suggested that *OsFIE1* evolved from *OsFIE2* by gene duplication after the tribe Oryzeae had evolved. Findings showed that *OsFIE1* and *OsFIE2* have evolved distinct functions in seed and endosperm development, and that they experienced different natural selection pressures.

## Results

### Divergence of the *FIEs* in rice

*OsFIE1* (Os08g0137100) and *OsFIE2* (Os08g0137250) are closely arrayed on chromosome 8 of rice (*Oryza sativa*), and are separated by a putative actin gene (**Supplementary Fig. 1A**). Such an arrangement is highly conserved among different *Oryza* species, such as *O. meridionalis* (A genome), *O. punctata* (B genome) and *O. brachyantha* (F genome) (**Supplementary Fig. 1A**), but differs from that in maize, where the two FIE homologs are located on different chromosomes (Danilevskaya *et al.*, 2003b). Next, we used the amino acid sequences of the OsFIEs as queries to search against the genomes of 20 diploid monocot species that had been deposited in the EnsemblPlants database, namely 11 members of the Oryzeae, four members of the Pooideae subfamily, three species of the Panicoideae subfamily and two non-grass species (*Dioscorea rotundata* and *Musa acuminata*). Most of the monocots (16/21) we studied have two *FIE* genes, while *Dioscorea rotundata, Musa acuminate, Setaria italica* and *Hordeum vulgare* have one and *Brachypodium distachyon* has four (**Fig. 1A**). Our phylogenetic analysis indicated that the duplication events of the *FIE*s probably occurred independently in different monocot genera. For example, two *FIE*s of maize were grouped together, but *OsFIE1* and *OsFIE2* belonged to two different clades (**Fig. 1A**). The *OsFIE1* homologs were exclusively found in the Oryzeae tribe and were relatively distant from the other *FIE* homologs (**Fig. 1A**). The *FIE2* homologs of the Oryzeae were closer to the *FIE*s of non-Oryzeae members of the other monocot species (**Fig. 1A**). The findings strongly suggested that *FIE1* of the Oryzeae evolved from *FIE2*, and that the duplication event probably occurred after the origin of the tribe Oryzeae.

**Figure 1.**
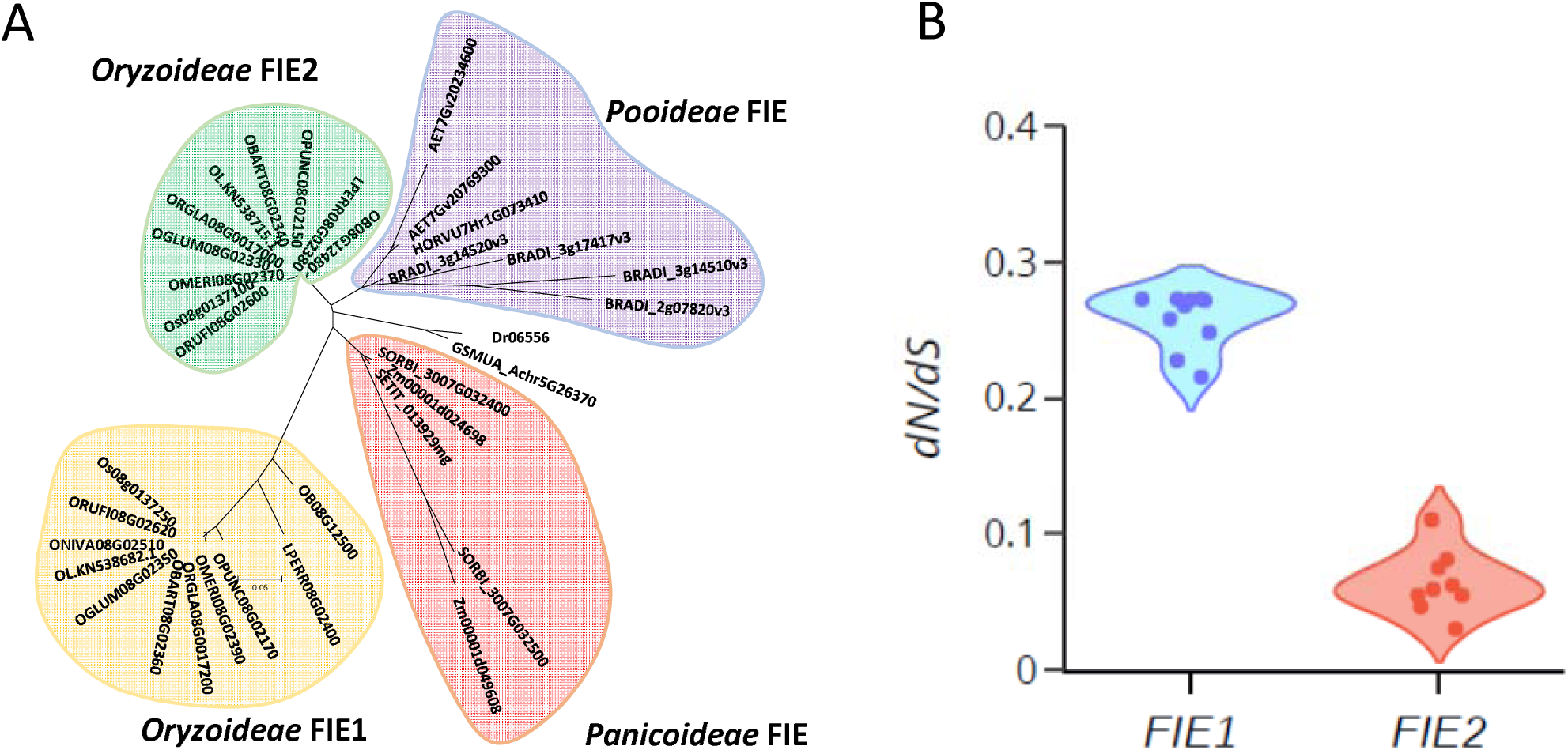
Evolutionary analysis of FIE homologs of rice. (A). Phylogenetic tree of the FIE homologs of monocots. EnsemblPlants IDs of the homologs were ovided. The initials of the accession numbers indicate the origin of the gene. All the rice homolog s start with “O”. LPRER, AET, HORVU, BRADI, SORBI, Zm, SETIT, GSMUA and Dr indicates *Leersia rrieri, Aegilops tauschii, Hordeum vulgare, Brachypodium distachyon, Sorghum bicolor, Zea mays, taria italica, Musa acuminata and Dioscorea rotundata*, respectively. The maximum likelihood ethod was used for the tree construction. (B). Violin plot of *dN*/*dS* of different *FIE1* and *FIE2* homologs in the genus *Oryza. L. perrieri* was used an out-group for the *dN*/*dS* calculations.

Compared with OsFIE2, OsFIE1 had an additional 82 amino acid (aa) residues at its N-terminal end (**Supplementary Fig. 1B**). This tail was conserved among *Oryza* species, but the FIE1 homolog of *Leersia perrieri*, which belongs to a different genus of the tribe Oryzeae, did not carry this tail (**Supplementary Fig. 2**). This indicated that this extra N-terminal tail of OsFIE1 had probably evolved after the genera *Oryza* and *Leersia* had diverged, which occurred 14.2 million years ago (Kellogg, 2009). Using *L. perrieri* as an outlier group, we calculated the ratio of the non-synonymous (*dN*) to synonymous (*dS*) substitutions of the *FIE1* and *FIE2* coding sequences of different *Oryza* species. The results showed that the *dN*/*dS* ratio of *FIE2* (0.03–0.11) was significantly lower than that of *FIE1* (0.21–0.27), suggesting that *FIE2* had experienced the more intense purifying selection in rice (**Fig. 1B**).

In line with previous findings (Li *et al.*, 2014), we found that *OsFIE1* and *OsFIE2* displayed quite different spatiotemporal expression patterns. Expression of *OsFIE1* was specifically activated in the endosperm, whereas *OsFIE2* is ubiquitously expressed (**Supplementary Fig. 3A and B**). *OsFIE1* and *OsFIE2* showed similar expression profiles during caryopsis development, being highly upregulated from 3 to 10 d after fertilization (DAF) (**Supplementary Fig. 3C**). However, *OsFIE2* transcripts were more abundant than those of *OsFIE1* in the developing caryopsis. The findings suggested that *OsFIE1* had probably been subfunctionalized as a result of functional divergence following duplication. Divergence of expression usually reflects a change in gene regulation. We had previous found that application of a DNA methylation inhibitor, 5-aza-2’-deoxycytidine, could ectopically activate expression of *OsFIE1* in seedlings, whereas *OsFIE2* show limited response to the chemical (Chen *et al.*, 2018a). We therefore proposed that *OsFIE1* and *OsFIE2* might have distinct DNA methylation patterns in the genome. To test this hypothesis, we performed bisulfite-PCR (BS-PCR) sequencing to detect DNA methylation of the two genes. Generally, *OsFIE2* showed higher levels of methylation level than *OsFIE1* in the promoter region as well as in the coding sequence (**Supplementary Fig. 3D**). Overall, the methylation level of *OsFIE2* was stable across leaf and endosperm tissues. However, *OsFIE1* was generally hypomethyalted in the endosperm at 6 DAF in comparison with the leaf (**Supplementary Fig. 1A**).

### Deleterious effects of ectopically expressed *OsFIE1*

Using the ubiquitin promoter of maize to drive the ectopic expression of *OsFIE1* (*Ubi*::*OsFIE1*) resulted in substantially decreased plant height and increased tiller numbers (**Fig. 2A and B**), phenotypic effects which were consistent with previous findings (Zhang *et al.*, 2012; Folsom *et al.*, 2014). We then analyzed histone modifications of the *Ubi*::*OsFIE1* lines using antibodies that react exclusively against tri-methylated histone H3 at lysine 27 (H3K27me3). The results showed that H3K27me3 was significantly elevated in *Ubi*::*OsFIE1* (Fig. 2C). This finding indicated that, similar to OsFIE2 (Nallamilli *et al.*, 2013), the OsFIE1-PRC2 complex has histone methyltransferase activity for H3K27me3 *in vivo*. We also used the *OsFIE2* promoter to drive the expression of *OsFIE1* (*pOsFIE2*::*OsFIE1*) in rice (**Fig. 2D and E**). The transgenic plants showed moderate (∼20-fold) up-regulation of *OsFIE1* expression in leaves (**Fig. 2E**). However, the *OsFIE2*::*OsFIE1* lines did not show the dwarf phenotype as was displayed by *Ubi*::*OsFIE1* (**Fig. 2A**). These findings suggested that the adverse effects of ectopically expressed *OsFIE1* occurred when *OsFIE1* products accumulated to a high concentration in vegetative tissues. However, moderate up-regulation, resembling that of *OsFIE2*, appeared not to induce these defects. Therefore, the adverse effects are dosage dependent.

**Figure 2.**
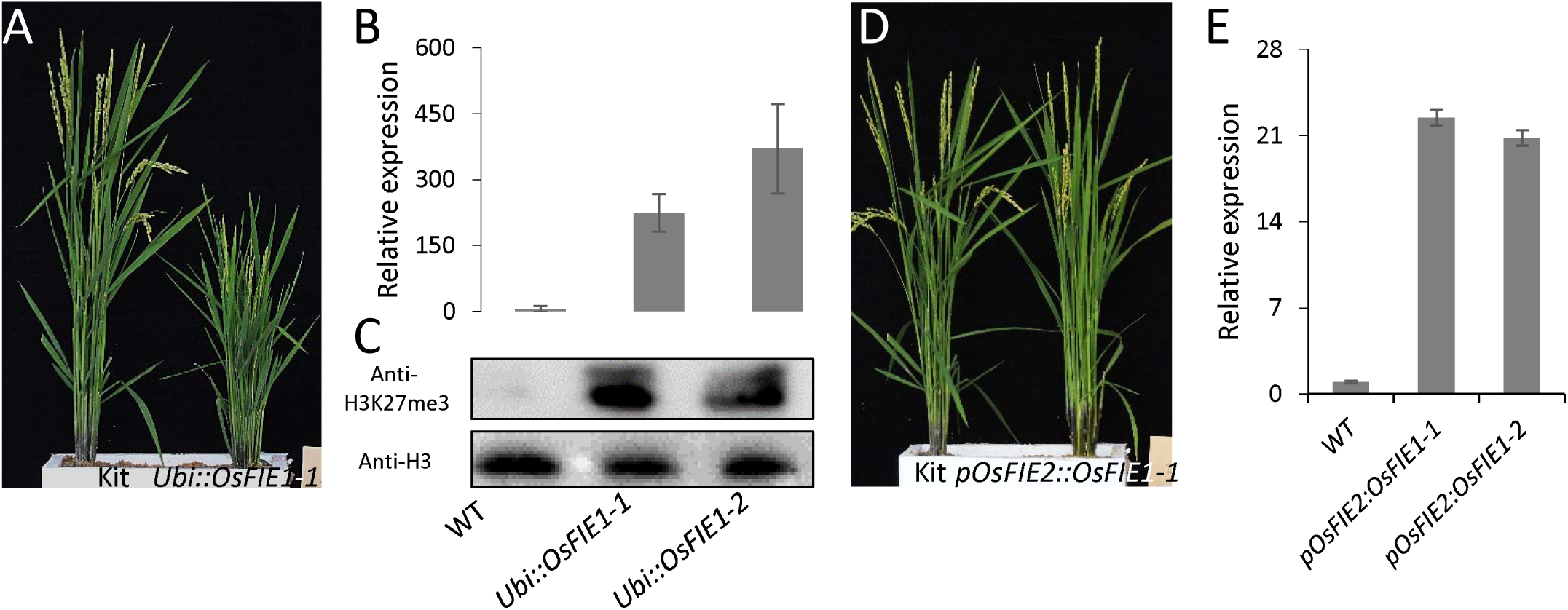
Deleterious effects of ectopically expressed *OsFIE1* on vegetative growth are dosage dependent. (A). Phenotype of the wild type (WT) and a representative *OsFIE1*-overexpression line (*Ubi::OsFIE1*) at the heading stage. (B). Relative expression of *OsFIE1* in leaves of *Ubi::OsFIE1* plants. (C). H3K27me3 was elevated in *Ubi::OsFIE1*. (D). Phenotype of WT and a *proOsFIE2::OsFIE1* line at the heading stage. (E). Relative expression of of *OsFIE1* in leaves of *pOsFIE2::OsFIE1* plants.

OsFIE1 has an extra tail segment at the N-terminus (**Supplementary Fig. 1B**). To discuss whether the tail contributes to the deleterious phenotypic effects on vegetative growth, we generated two chimeric OsFIEs, namely Chi-OsFIEa and Chi-OsFIEb, by swapping the N-terminus of OsFIE1 to OsFIE2 as illustrated in **Fig. 3A**. Overexpression of the chimeric *OsFIEs* driven by the ubiquitin promoter of maize (**Fig. 3B**) did not cause growth defects as had been displayed in *Ubi*::*OsFIE1* (**Fig. 3C and D**). The plant height and tiller number of *Chi-OsFIEa* and *Chi-OsFIEb* were comparable to the corresponding values for the wild type (WT) but were significantly different from those in *Ubi*::*OsFIE1* (**Fig. 3E and F**). The results suggested that the extra N-terminal segment of OsFIE1 alone was not the cause of the vegetative defects observed.

**Figure 3.**
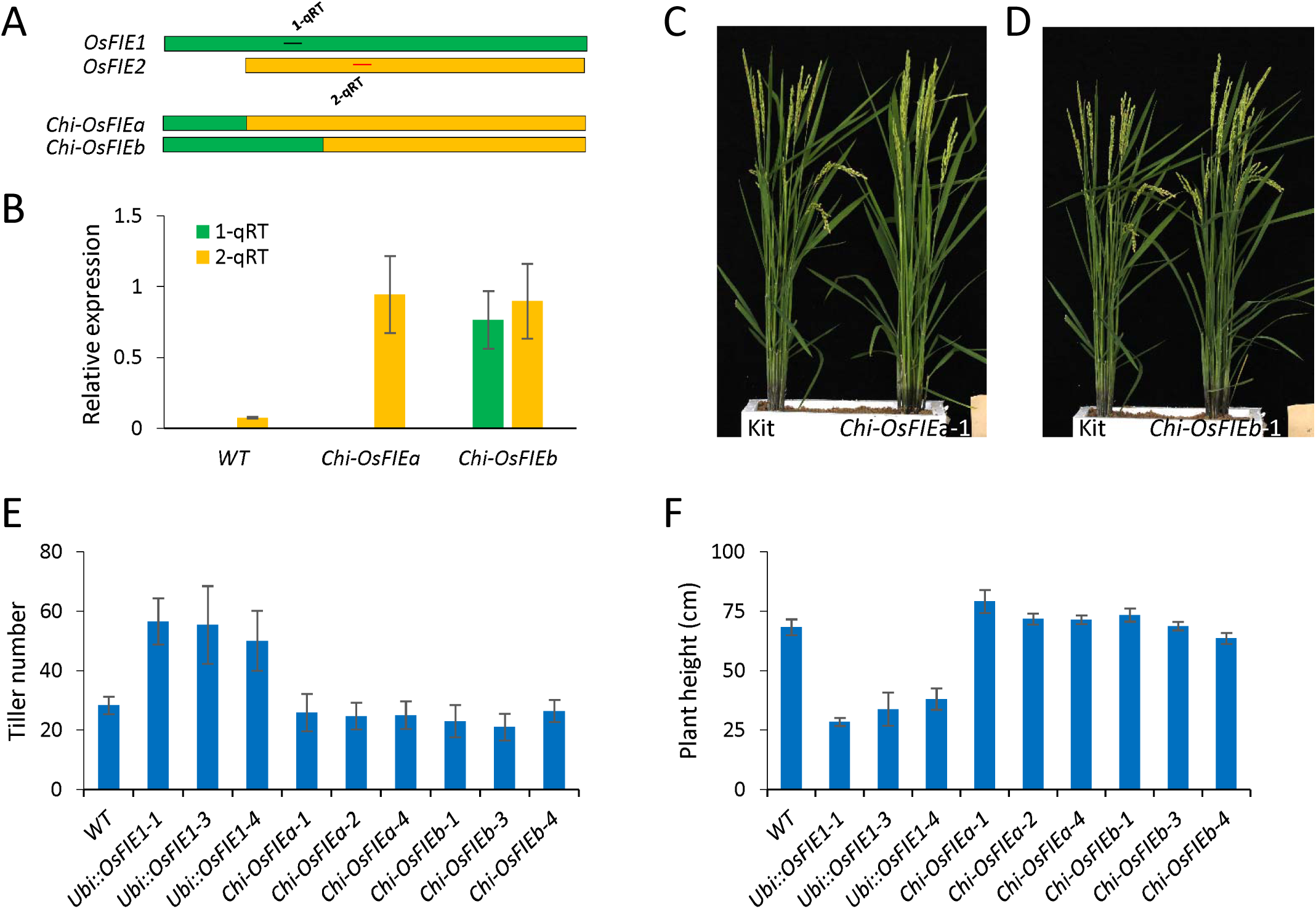
Ectopic expression of the chimeric *OsFIE* did not result in vegetative defects. (A). Scheme of the chimeric OsFIEs, achieved by swapping the extra N-terminal tail of OsFIE1 to IE2. The green and yellow bars indicated OsFIE1 and OsFIE2 proteins, respectively. (B). Confirmation of ectopic expression of the chimeric *OsFIEa* (*Chi-OsFIEa*) and *Chi-OsFIEb* in sgenic plants. 1-qRT and 2-qRT indicated the segments used to distinguish the *OsFIE1*- and *OsFIE2*-origin transcripts. The corresponding positions of 1-qRT1 and 2-qRT are indicated in (A). Three ogical replicates were used; the error bars indicated standard deviations. (C, D). Phenotypes of *Chi-OsFIEa-1* (C) line and *Chi-OsFIEb-1* (D) plants at the heading stage. (E, F). Tiller numbers (E) and plant height (F) of different transgenic lines of *Ubi::OsFIE1, Chi-OsFIEa Chi-OsFIEb*. Twenty plants for each line were measured; error bars indicate standard deviations.

### Up-regulation of *OsFIE1* expression in endosperm reduced the seed size of rice

Previous studies had suggested that overexpression of *OsFIE1* led to reduced seed size (**Fig. 4A**; Zhang *et al.*, 2012; Folsom *et al.*, 2014). However, it was unclear whether the phenotype was directly caused by the overexpression of *OsFIE1* in seeds, or whether it was an indirect effect of ectopic expression of *OsFIE1*, reflecting the reduced stature and impaired vegetative growth (**Fig. 2A**). To clarify the effects of *OsFIE1* overexpression on seed development, we specifically activated its expression in the endosperm, using the promoter of the *rice glutelin 1* gene (Russell & Fromm, 1997) to drive *OsFIE1* (*GT1*::*OsFIE1*). The expression of *OsFIE1* in the endosperm in independent transgenic lines was significantly increased, whereas the expression of *OsFIE2* was not altered in the transgenic plants (**Fig. 4B**). Unlike the *Ubi*::*OsFIE1* plants, the *GT1*::*OsFIE1* lines showed vegetative development similar to that of the WT (**Fig. 4C**). However, all three independent *GT1*::*OsFIE1* transgenic lines analyzed produced seeds significantly smaller than the WT seeds (**Fig. 4A and D-F**). One-thousand-grain weight of *GT1*::*OsFIE1* was reduced to 86–88% of the WT value (**Fig. 4G**). The results unambiguously indicated that overexpression of *OsFIE1* could directly affect seed development.

**Figure 4.**
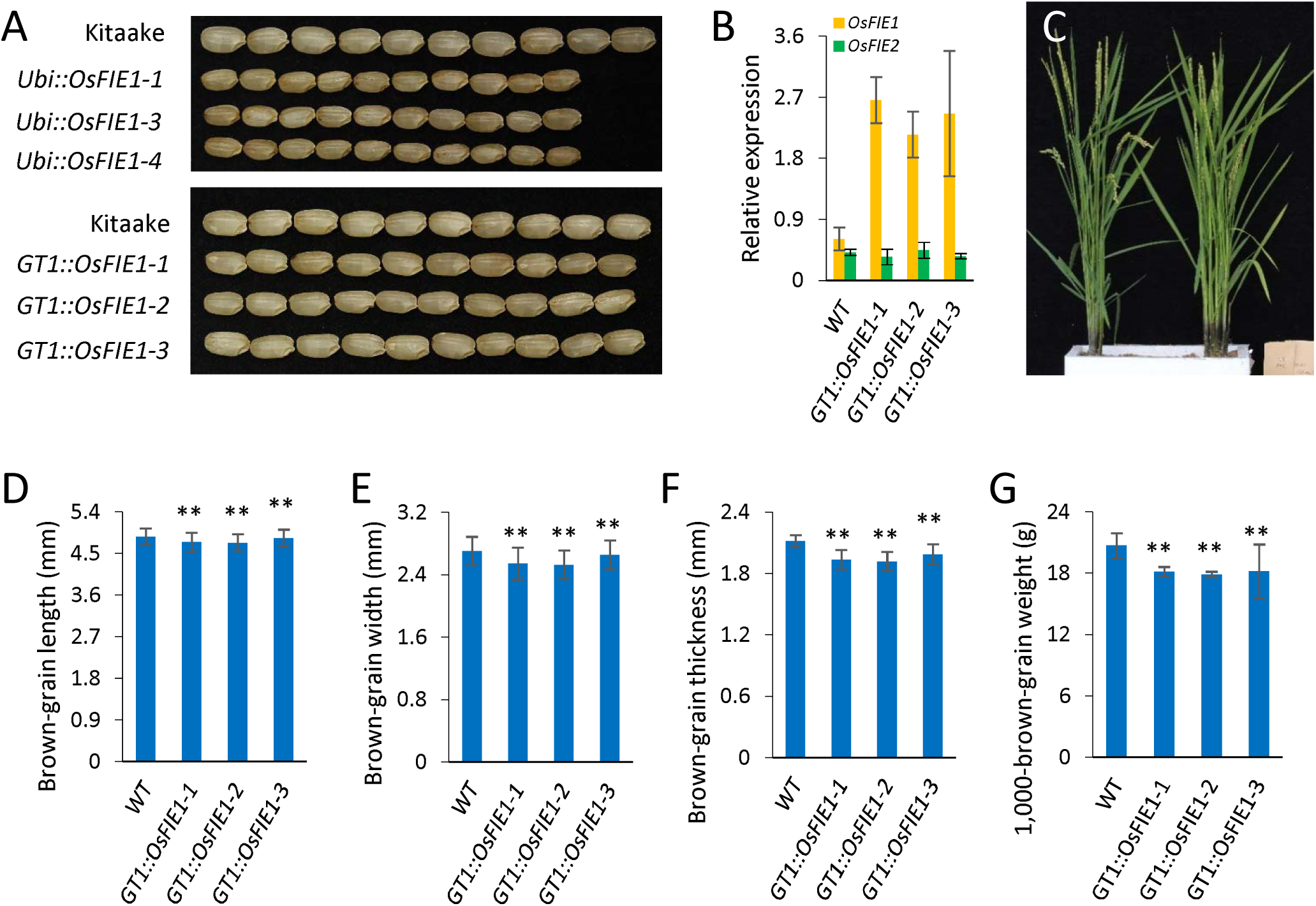
Overexpression of *OsFIE1* resulted in reduced seed size. (A). Phenotypes of seeds from *Ubi::OsFIE1* and *GT::OsFIE1*. (B). Confirmation of the expression up-regulation of *OsFIE1*, but not *OsFIE2*, in the caryopses (6 days r fertilization (DAF)) of different *GT1::OsFIE1* transgenic lines. (C). Phenotypes of the wild-type (WT) and *GT::OsFIE1* plants at the heading stage. (D-G). Length (D), width (E), thickness (F) and 1000-grain weight (G) of the brown seeds produced by *GT1::OsFIE1* lines.

### Mutant *osfie1* showed smaller seeds and increased pre-harvest sprouting

To investigate the function of *OsFIE1*, we generated two independent CRISPR/Cas9 *osfie1* mutant lines using different targets (1-T1 and 1-T2) (**Fig. 5A**). The vegetative growth of *osfie1* was not distinguishable from that of the WT (**Fig. 5B**). In line with the findings by Huang *et al.* (2016), our results showed that the seed size and seed weight of *osfie1* were greatly reduced relative to the WT (**Fig. 5C and D**), these effects being associated with reduced seed width and thickness, rather than shorter seed length (**Fig. 5E-G**).

**Figure 5.**
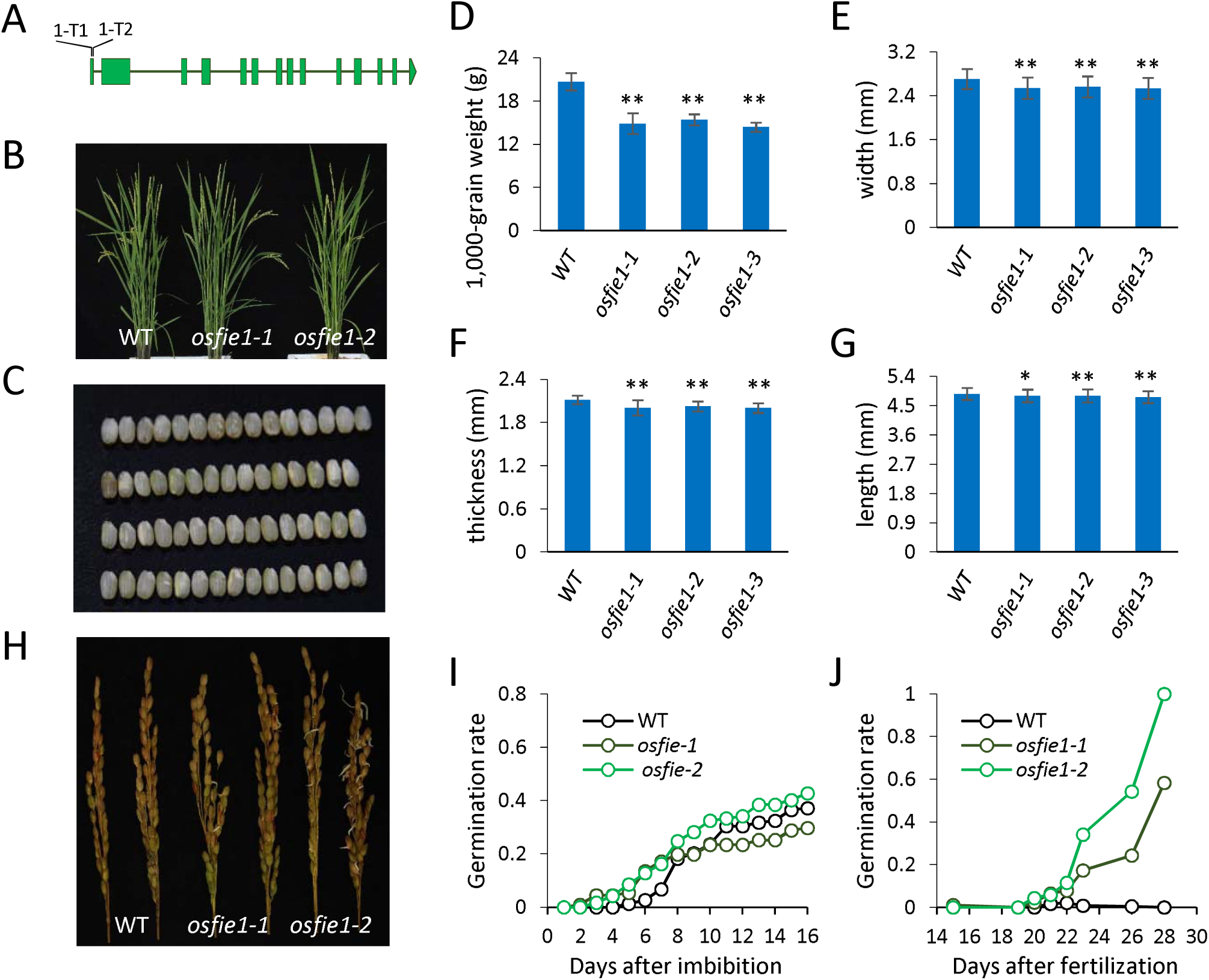
Phenotyping of the osfie1 mutants. (A). Illustration of the targets (1-T1 and 1-T2) that were used for generation of CRISPR/Cas9 mutants. (B). Phenotype of a wild type (WT) plant and two osfie1 mutants at the heading stage. (C). Seed phenotype of WT and *osfie1* mutants. Images from top to bottom are WT, *osfie1-1, osfie1-2* and *osfie1-3*, respectively. (D, G). 1000-grain weight (D), length (E), width (F), and thickness (G) of the brown seeds produced by *osfie1* lines. (H). Reduced dormancy of the *osfie1* mutants. (I). Dynamic curves of germination of WT, *osfie1-1* and *osfie1-2*. (J). Germination rate of different-aged seeds of WT, *osfie1-1* and *osfie1-2*.

Notably, we observed that *osfie1* exhibited a significantly higher pre-harvest sprouting phenotype in the paddy field in each of two seasons (**Fig. 5H**), which suggested that *OsFIE1* might positively regulate dormancy in rice. To confirm this, we carried out germination tests on near-mature seeds (∼30 DAF) that had been directly collected from field without drying. More than 15 % of the seeds of *osfie1-1* and *osfie1-2* geminated, but less than 7% WT seeds geminated after 7 d after imbibition (DAI) (**Fig. 5I**). Next, we used differently aged seeds of different genotypes in the germination test (**Fig. 5J**). The results showed that *osife1* had a significantly higher germination potential than WT in terms of the immature seeds, with more than 20% of the 23 DAF seeds germinating at 7 DAI, which was significantly higher than that of the WT (**Fig. 5J**).

### *OsFIE2*, but not *OsFIE1*, is required for cellularization of the endosperm

Previous studies had shown that *OsFIE2* RNAi plants had a lower seed-set rate, with some of the seeds produced being abnormal (Nallamilli *et al.*, 2013; Li *et al.*, 2014), but the mechanism by which *OsFIE2* affects seed development is still largely unknown. Because of the strong similarities between *OsFIE1* and *OsFIE2* (**Supplementary Fig. 1B**), we believed that it would not be a good strategy to use RNAi lines for the study of seed or endosperm development, owing to the overlap in terms of spatiotemporal expression of these two genes in seeds (**Supplementary Fig. 1E**). Therefore, we made efforts to generate *osfie2* mutants using a CRISPR/Cas9 approach (**Fig. 6A**). Unfortunately, despite the use of three different targets (2-T1, 2-T2 and 2-T3), our attempts to obtain callus-regenerated homozygous mutants of *OsFIE2* at the T_0_ generation failed, although heterozygous *OsFIE2*^+*-*^ plants were readily obtained (**Fig. 6B**). In contrast, we could obtain homozygous mutants of *OsFIE1* at a high frequency (∼30%) in the T_0_ generation (**Fig. 6B**). In addition, we used a single construct with a cassette consisting of both 1-T1 and 2-T2, attempting to knock-out *OsFIE1* and *OsFIE2* simultaneously. We succeeded in obtaining the *osfie1* mutant in the homozygous condition at the *OsFIE1* locus, but again only heterozygous (*osfie1*/*OsFIE2*^+*-*^) were obtained at the *OsFIE2* locus (**Fig. 6B**). According to these findings, we inferred that *OsFIE2*, but not *OsFIE1*, is indispensable for rice regeneration. Alternatively, *OsFIE2* is more essential for development, because the transformants lacking functional *OsFIE2* were not viable.

**Figure 6.**
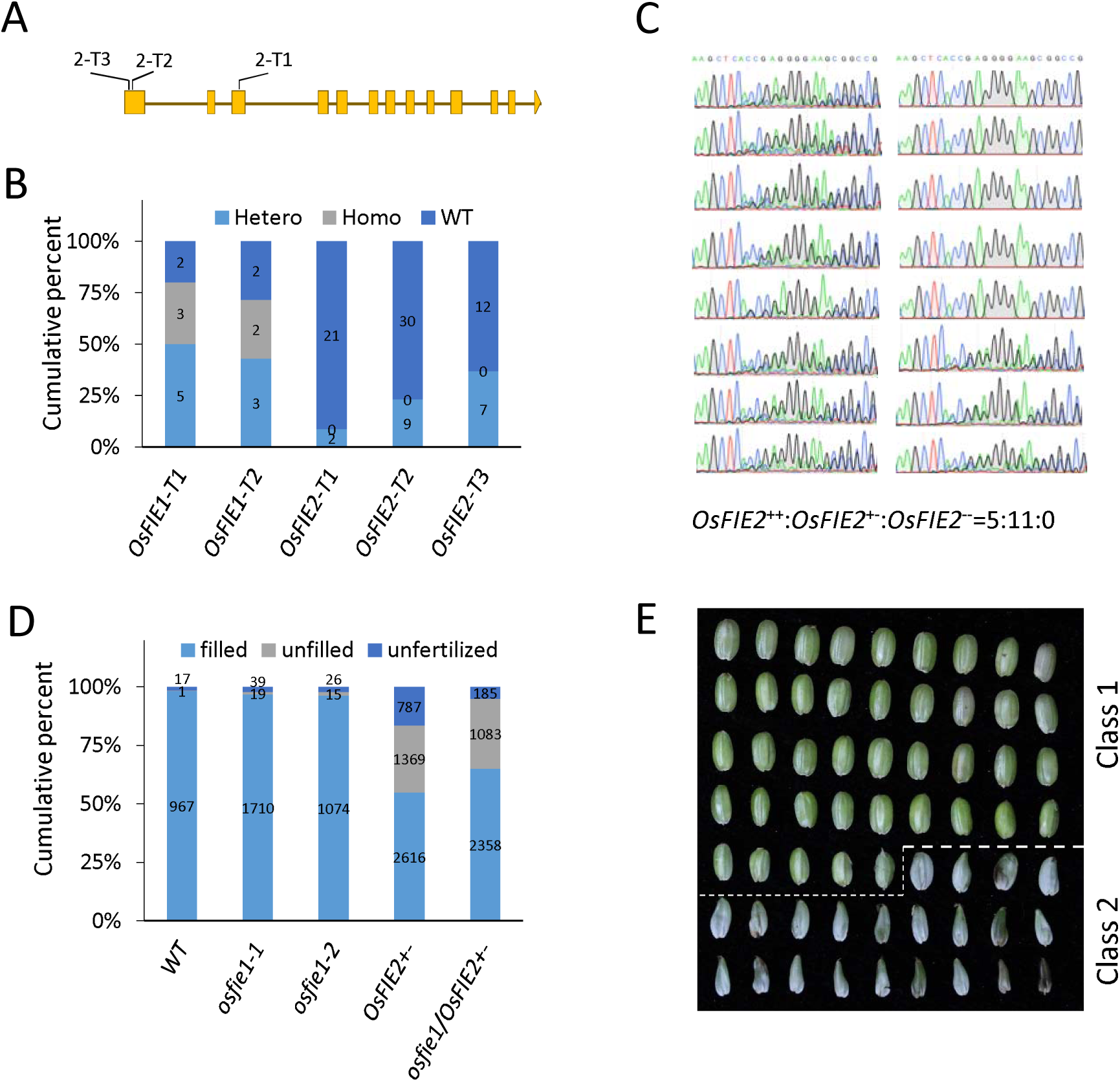
osfie2 and *osfie1,2* mutants were leathal. (A). Schematic drawing of the three targets (2-T1, 2-T2 and 2-T3) that were used for generation of PR/Cas9 mutants of *osfie2*. (B). Cumulative percentage of the not-edited (wild type, WT), monoallele-edited (Hetero) and diallele-edited homozygous (Homo) individuals that were regenerated from *Agrobacterium*-mediated transformation at T_0_ generation. The number of individuals of each genotype are indicated on the bars. (C). Sequencing of 16 F_2_ individuals derived from *OsFIE2*^+−^. (D). Cumulative percentage of well-filled, unfilled and unfertilized seeds produced by WT (n = 985), *osfie1-1* (n = 1768), *osfie1-2* (n = 1115), *OsFIE2*^+−^ (n = 4772) and *osfie1*/*OsFIE2*^+−^ (n = 3626). (H). Caryopses collected from a single panicle of an *OsFIE2*^+−^ plant at 15 days after fertilization (DAF). Two classes of caryopsis were observed: Class 1 that produced starchy endosperm, and Class 2 that duced watery endosperm.

In a segregating population from self-pollinated *OsFIE2*^+−^, only *OsFIE2*^++^ and *OsFIE2*^+−^ plants were obtained (**Fig. 6C**). After 5 DAF, the seeds produced by *OsFIE2*^+-^ could readily be categorized into two seed classes (**Fig. 6E**). Seeds of Class 1 showed an appearance similar to that of the WT, while seeds of Class 2 showed growth cessation frequently, illustrated with a swollen belly consisted watery endosperm (**Fig. 6E**). At maturity, the Class 2 seeds were completely empty (**Supplementary Fig. 4**). In agreement with this observation, the seed-set rate of *OsFIE2*^+-^ and *osfie1*/*OsFIE2*^+-^ was significantly lower than that of WT or *osfie1* (**Fig. 6D**). Notably, the rate of unfertilized seeds, which failed to show enlargement of the caryopses, was significantly increased in *OsFIE2*^+-^ (16.5%) and *osfie1*/*OsFIE2*^+-^ (5.1%) in comparison to that of the WT (1.7%). This was likely not caused by lower viability of the male or female gametes. Because the development of pollens and embryo sacs seemed normal in *OsFIE2* and *osfie1*/*OsFIE2* (**Supplementary Fig. 5A-H**).

The Class 1 seeds produced a starchy endosperm at 15 DAF, whereas the Class 2 seeds produced a watery endosperm instead at this time point (**Fig. 6E**). Because we had failed to obtain the *osfie2* mutant, we believed that the Class 2 seeds were the homozygotes. The Class 1 seeds were morphologically different from the Class 2 seeds after 4–5 DAF. Early endosperm development of Class 2 seeds was impaired (**Fig. 7A-L**). In seeds of WT and *osfie1* at 4 DAF, cellularized endosperm cells completely filled the embryo sac (**Fig. 7I and J**), but only a few layers of endosperm cells were observed in *osfie2* or *osfie1,2* caryopses at this time point (**Fig. 7K and L**), which indicated that cellularization of the Class 2 seeds was delayed or impaired. At 7 DAF, most of the *osfie2* seeds still had an empty vacuole. A few seeds had multiple layers of loosely packed endosperm cells (**Fig. 7M and N**), but much less starch had accumulated in these cells, as revealed by I_2_-KI staining assay, compared with WT seeds (**Fig. 7O and P**). In line with this observation, expression of some key genes involving starch biosynthesis, including *OsAGPL2, OsPUL, OsGBSSI, OsISA2, RPBF* and *RISBZ1*, were substantially down-regulated in *osfie2* and *osfie1,2* seeds at 6 DAF (**Fig. 7Q-V**). On the other hand, expression of *OsAPGL1* and a negative regulator gene of starch biosynthesis, *RSR1*, were upregulated (**Fig. 7W and X**). The I_2_-KI staining experiment showed that starch accumulation was not affected in the seed coat of *osfie2* (**Fig. 7O and P**), suggesting that the development of the seed coat and integument were probably not strongly impaired. Notably, early endosperm development of *osfie1* seemed not to be affected (**Fig. 7B, F and J**). Together, the results suggested that *OsFIE2*, but not *OsFIE1*, is essential for early endosperm development in rice.

**Figure 7.**
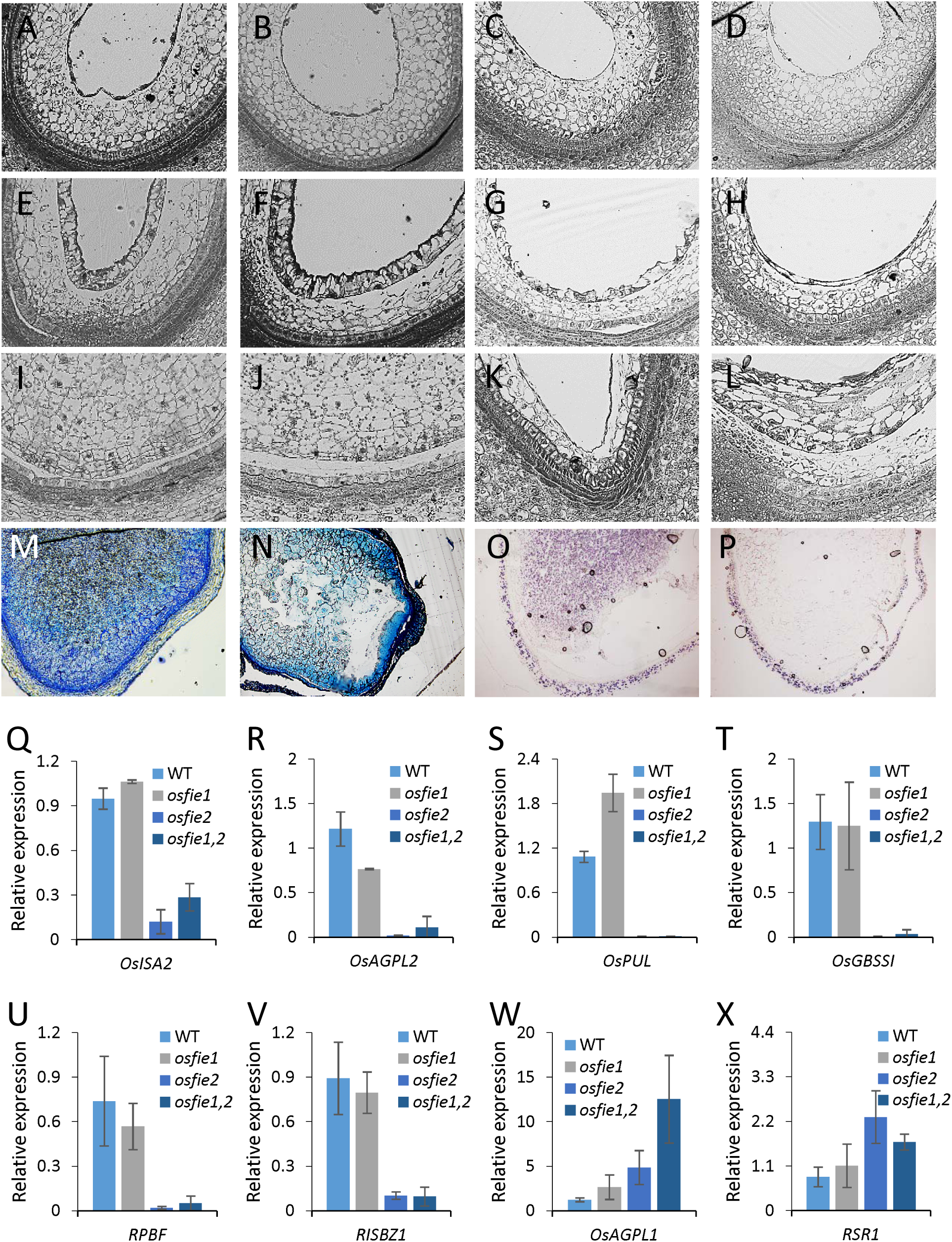
Early endosperm development of wild type (WT), *osfie1, osfie2* and *osfie1,2*. (A-L). Sections of 2 days after fertilization (DAF) (A-D), 3 DAF (E-H) and 4 DAF (I-L) endosperm of WT (A, E, I), *osfie1-1* (B, F, J), *osfie2* (C, G, K) and *osfie1,2* (D, H, L). (M, N). Sections of 7 DAF endosperm of WT (M) and *osfie2* (N). (O, P). I_2_-KI staining of 7 DAF endosperm of WT (O) and *osfie2* (P). (Q, X). Expression of some key genes involved in starch biosynthesis. Three biological replicates were used; error bars indicated standard deviations.

The embryo development of neither *osfie2* nor *osfie1,2* was significantly affected at 4 DAF, relative to the WT (**Supplementary Fig. 6**). As with the WT, globular embryos could be observed in the mutants at 4 DAF (**Supplementary Fig. 6A-C**). However, embryos of *osfie2* were not still observable at 15 DAF (**Supplementary Fig. 6D and E**). The embryos had probably degenerated by this point, leaving a cavity in most of the seeds which could be seen (**Supplementary Fig. 6D and E**). We found embryos in two of the five *osfie1,2* seeds we studied, but the development of these had already ceased (**Supplementary Fig. 6F**). The results suggested that the embryogenesis of *osfie2* and *osfie1,2* was arrested, as was endosperm development.

### Autonomous endosperm production was occasionally observed in *osfie2* mutants

Mutation of *FIE* in Arabidopsis can lead to autonomous endosperm production without fertilization (Ohad *et al.*, 1996; Chaudhury *et al.*, 1997; Kiyosue *et al.*, 1999). However, whether *OsFIE1* and *OsFIE2* play similar roles in the repression of ovary proliferation in rice is not clear. In the present study, WT, *osfie1, OsFIE2*^+*-*^ and *osfie1*/*OsFIE2*^+*-*^ panicles were emasculated before flowering and bagged to prevent cross-pollination. No autonomous seeds were produced by the 162 emasculated spikelets of WT and the 76 emasculated spikelets of *osfie1* (**Fig. 8A**). However, *OsFIE2*^+-^ and *osfie1*/*OsFIE2*^+-^ were able to produce autonomous seeds at a very low frequency (2.6% of *OsFIE2*^+-^ and 4.9% of *osfie1*/*OsFIE2*^+-^) (**Fig. 8A**). The autonomous seeds were not able to accumulate starch. They were completely empty when dried (**Fig. 8B and C**).

**Figure 8.**
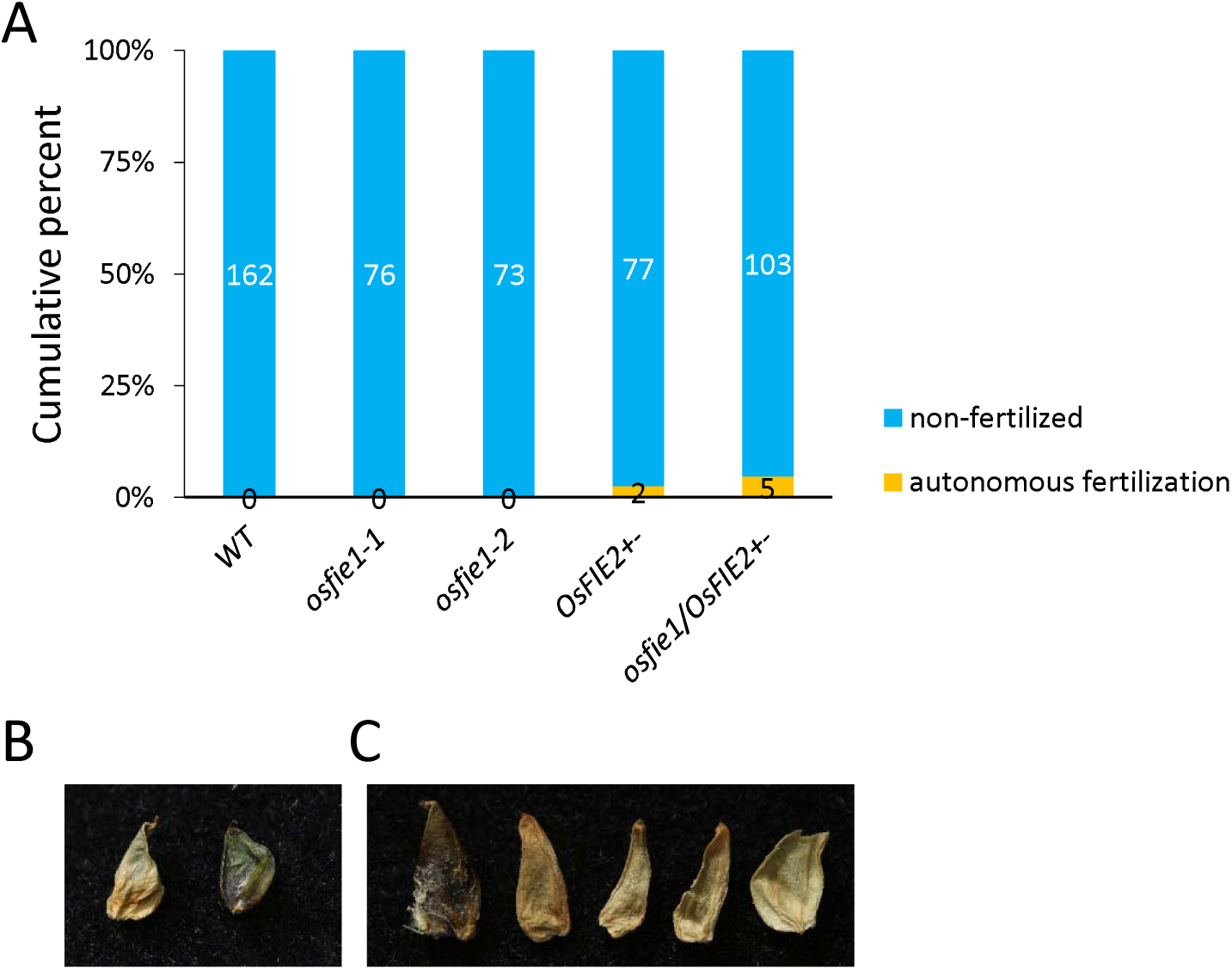
Autonomous fertilization of *OsFIE2*^+−^ and *osfie1*/*OsFIE2* ^+−^. (A). Cumulative percentages of the non-fertilized and autonomously fertilized seeds in wild type (WT), *osfie1-1, osfie1-2, OsFIE2*^+−^ and *osfie1*/*OsFIE2*^+−^. The number of each type of seed was indicated on the bars. (B, C). Morphology of dried autonomous seeds of *OsFIE2*^+−^ (B) and *osfie1*/*OsFIE2*^+−^ (C).

### Additive effects of *OsFIE1* and *OsFIE2* on gene expression

In order to dissect how *OsFIE1* and *OsFIE2* function in seed development, we carried out RNA-Seq to discover differentially expressed genes (DEGs) from *osfie1, osfie2* and the *osfie1,2* double mutant. In caryopses at 5 DAF, using the cutoff thresholds of FC (fold change) >2 and FDR (false discovery rate) <0.05, 112 and 701 genes were down- and upregulated, respectively, in *osfie1* when compared with WT (**Fig. 9A and B**). The number of DEGs was much higher in *osfie2* and *osfie1,2*. A total of 1,110 down-regulated DEGs and 6,175 upregulated DEGs were identified from *osfie2*, whereas 1,152 down-regulated DEGs and 5,826 upregulated DEGs were identified from *osfie1,2* (**Fig. 9A and B**). Most of the DEGs were upregulated relative to the WT (**Fig. 9A and B**), a finding which was consistent with the functions of OsFIE1 and OsFIE2, which are involved in H3K27me3 modification of chromatin.

**Figure 9.**
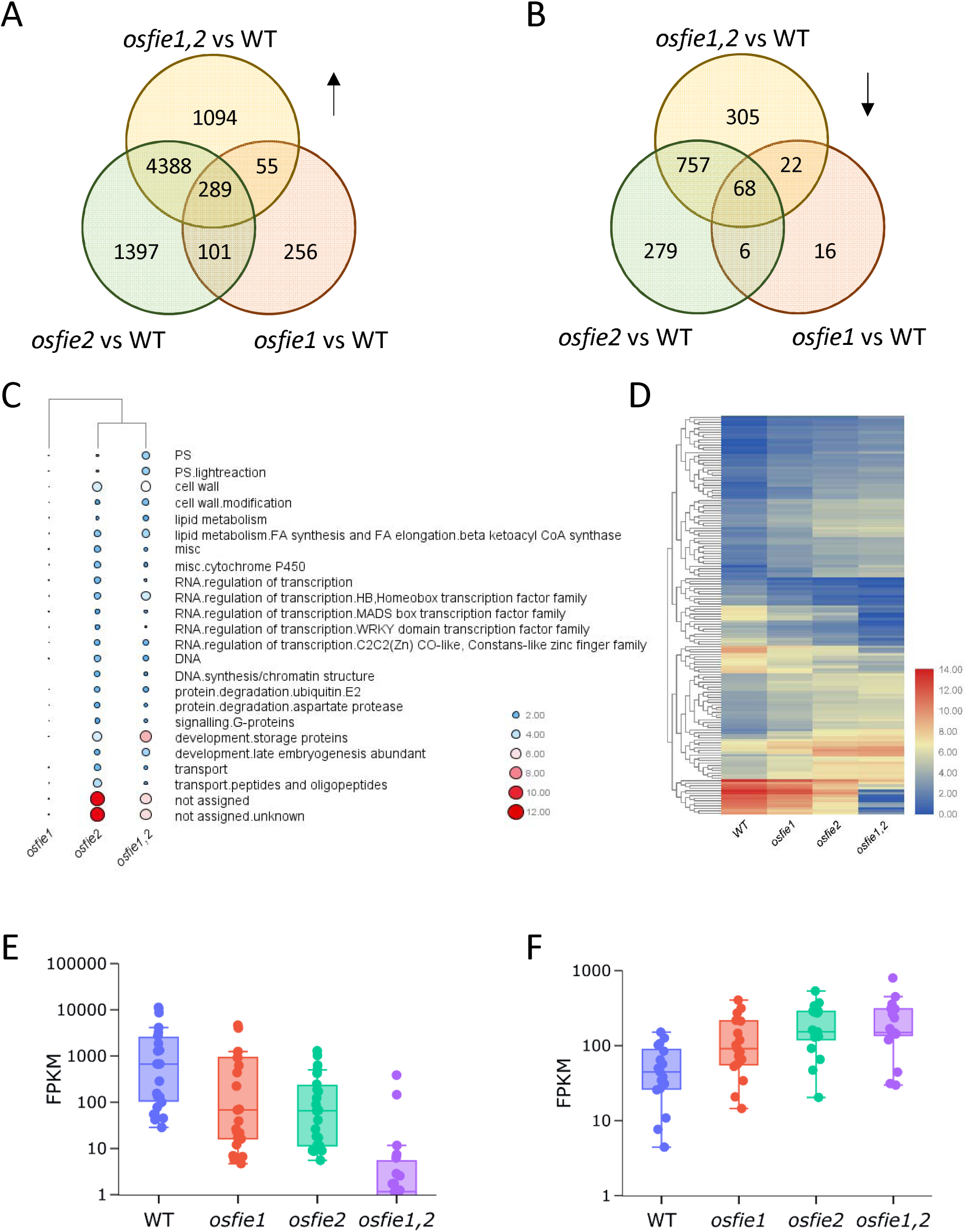
Transcriptome analysis of caryopses 5 days after fertilization (5 DAF) of wild type WT, *osfie1, osfie2* and *osfie1,2*. (A, B). Venn diagrams of the upregulated (A) and down-regulated (B) genes identified from *osfie1, osfie2* and *osfie1,2* in comparison to WT. (C). MapMan pathway enrichment analysis of the differentially expressed genes (DEGs). Circle size and colors indicate the log scale of the enrichment. (D). Heatmap of the expression of DEGs common to *osfie1, osfie2* and *osfie1,2*. Gene expression was indicated by log2(FPKM, Fragments Per Kilobase of transcript per Million fragments mapped). (E, F) Additive effects of *OsFIE1* and *OsFIE2* on expression of storage protein biosynthesis-related (E) genes and photosynthesis-related genes (F).

Most of the DEGs identified from *osfie2* and *osfie1,2* overlapped (**Fig. 9A and B**). MapMan analysis suggested that genes related to storage protein biosynthesis, lipid metabolism, cell wall synthesis, and DNA synthesis were enriched (Benjamini-Hochberg corrected FDR < 0.05) in *osfie2* and *osfie1,2* (**Fig. 9C**). This result was in line with the observations that cellularization of *osfie2* was delayed and starch was not accumulated in *osfie2* endosperm (**Fig. 6**). Notably, several groups of transcription factor, such as homeobox, MADS box, WRKY and C2H2 CO-like zinc finger genes, were also enriched among the DEGs (**Fig. 9C**), indicating that these transcription factors play essential roles in early seed development of rice. Several type I MADS-box genes of Arabidopsis were found to act as negative regulators of cellularization. Notably, many MADS-box genes in rice, including MIKC-type and type I genes were significantly upregulated in *osfie1* and *osfie1,2* seeds (**Supplementary Fig. 7**). These genes may also be involved in the early endosperm development of rice.

Compared with *osfie2* and the *osfie1,2* double mutant, *osfie1* showed fewer effects on expression in seeds early in development (**Fig. 9A and B**), which further supported our hypothesis that *OsFIE1* is less important than *OsFIE2* for early seed development. Gene Ontology analysis revealed that genes involved in photosynthesis (FDR = 9.4e-05), cell wall organization or biogenesis (FDR= 9.4e-05) and response to stimuli (FDR= 0.0032) were significantly enriched among the DEGs identified from *osfie1*. About 85.7% of the down-regulated DEGs and 63.5% of the upregulated DEGs identified from *osfie1* also showed differential expression in *osfie2* or *osfie1,2* (**Fig. 9A and B**), suggesting that these genes are probably common targets regulated by OsFIE1 and OsFIE2.

Intriguingly, we found that the DEGs common to *osfie1, osfie2* and *osfie1,2* (**Fig. 9A and B**) were additively regulated by *OsFIE1* and *OsFIE2*. Overall, mutation of *OsFIE2* led to a larger effect than mutation of *OsFIE1*, whereas the *osfie1,2* double mutant showed the greatest change in expression (**Fig. 9D**). For example, among the common down-regulated DEGs (68 in total), twenty-three were storage protein-precursor-encoding genes. These genes were highly expressed in the WT, but were greatly suppressed in the mutants, with the degree of suppression being in the order *osfie1*<*osfie2*<*osfie1,2* (**Fig. 9E**). In contrast, 17 genes related to photosynthesis were upregulated in the mutant lines, which overall showed greatest transcript abundance in *osfie1,2* and moderate up-regulation in *osfie2*, while *osfie1* showed the least up-regulation (**Fig. 9F**). Taken together, these findings suggested that *OsFIE1* plays only a limited function in early seed development. However, *OsFIE1* can coordinate with *OSFIE2* in an additive manner to regulate most of the DEGs that were identified from *osfie1*.

## Discussion

The components of PRC2 are conserved between animals and plants. For animals, there are usually only one or two members of each of the PRC2 groups p55, E(z), Su(z)12 and ESC. However, plants usually have multiples of each group (Furihata *et al.*, 2016). For example, most of the monocots we analyzed in the present study had two or more *FIE* genes (**Fig. 1A**). The *FIE*s of the different graminaceous genera probably evolved as a result of independent duplication events. A duplication that occurred after the origin of the tribe Oryzeae generated two FIE orthologs, *OsFIE1* and *OsFIE2*, in rice. The phylogenetic and alignment analyses suggested that *OsFIE2* is the more ancient gene (**Fig. 1A and Supplementary Fig. 2**), because it is more similar to the *FIE*s of other grass species. *OsFIE2* could promote cellularization like *AtFIE* acts in Arabidopsis (Vinkenoog *et al.*, 2000), but *OsFIE1* showed only limited effects in early endosperm development in rice (**Fig. 7A-P**). The expression pattern and the methylation landscape of *OsFIE1* and *OsFIE2* were quite distinct (**Supplementary Fig. 1**), which suggested that the genes had probably diverged since they evolved. *OsFIE2* is universally expressed, but it is highly methylated in the promoter region when compared with *OsFIE1* (**Supplementary Fig. 2B and D**). Usually methylation act as a repression mark for gene expression (Deng *et al.*, 2016), whereas there have been findings suggesting that hypermethylation may also be involved in up-regulation of certain genes in crops (Lang *et al.*, 2017). Expression of *OsFIE1* is restricted to the endosperm (**Supplementary Fig. 2A**), possibly due to its deleterious effects when ectopically expressed in vegetative tissues (**Fig. 2A**). Previous studies had found that overexpression of *OsFIE1*, but not *OsFIE2*, could lead to dwarfism (Zhang *et al.*, 2012; Folsom *et al.*, 2014). In the current study, we found that the adverse effects caused by *OsFIE1* were dosage dependent (**Fig. 2**), with moderate up-regulation of *OsFIE1* in the seedling not resulting in developmental defects. So far, we still do not know why *OsFIE1*, but not OsFIE2, is harmful to vegetative growth. The “gene swapping” experiments suggested that the extra N-terminal tail of OsFIE1 only was not able to induce the deleterious effects (**Fig. 3**). The alignment analysis indicated that, in addition to the N-terminal difference, OsFIE1 has an extra 8-aa segment at the C-terminal end (**Supplementary Fig. 1B**). This difference, as well as the amino acid substitutions found in OsFIE1, may contribute to the deleterious effects.

The *osfie1* mutant produced smaller seeds and showed increased frequency of pre-harvest sprouting relative to the WT (**Fig. 5**), suggesting that *OsFIE1* plays roles in late seed development (**Fig. 10**). In support of this hypothesis, many DEGs identified in *osfie1* were responsible for storage protein biosynthesis (**Fig. 9C**). Interestingly, OsEMF2b, a member of Su(z)12 group in rice, has recently been found to function in seed dormancy by modulating the expression of *OsVP1* (Chen *et al.*, 2017). Therefore, OsFIE1 and OsEMF2b are possibly recruited into the same PRC2 that regulates seed dormancy and germination. In agreement, many regulators of late seed development and germination of Arabidopsis, such as *LEAFY COTYLEDON 2* (*LEC2*), *ABSCISIC ACID INSENSITIVE 3* (*ABI3*), *FUSCA 3* (*FUS3*), *ABI4* and *DELAY OF GERMINATION 1* (*DOG1*), were upregulated in *fie* mutant seedlings, possibly due to depletion of H3K27me3 of these genes (Bouyer *et al.*, 2011). The mechanism by which *OsFIE1* regulates dormancy in rice need to be further investigated. Because we failed to obtain an *osfie2* mutant, it is unclear whether *OsFIE2* functions on dormancy as well. However, PRC2-regulated seed dormancy and germination have been reported in monocots and dicots, suggesting that this function of PRC2 is conserved in plants. Taking into account that *OsFIE2* is probably the more ancient ortholog, we believe that *OsFIE2* functions in late seed development as well (**Fig. 10**).

**Figure 10.**
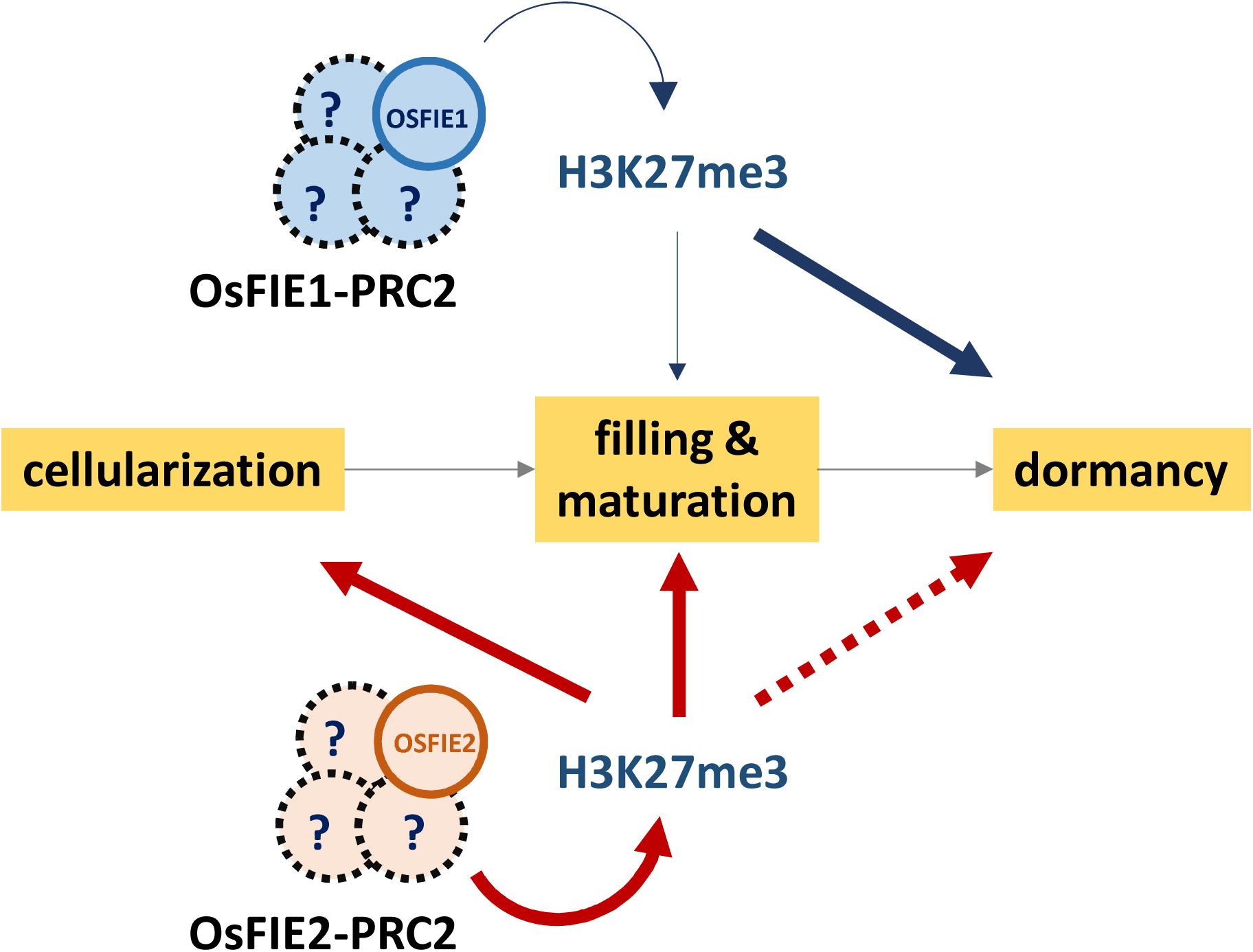
Functional divergence between *OsFIE1* and *OsFIE2* with respect to seed development of rice. Both OsFIE1 and OsFIE2 are able to interact with other members to form functional OsFIE1- and OsFIE2-PRC2. Whether the same or distinct components are recruited by OsFIE1 and OsFIE2 to form a polycomb complex is not determined. By modulating the H3K27me3 of its target genes, OsFIE2 acts as a positive regulator of endosperm cellularization and may also function in terms of starch filling of seeds. *OsFIE1* is less active on the regulation of cellularization, seed filling and maturation; but is essential for dormancy. It is not clear whether *OsFIE2* functions on dormancy as well. Dashed lines indicated undetermined components or regulation, and line thickness indicated importance for regulation.

Evidence from the present study suggested that *OsFIE2* is essential for early endosperm development *via* the promotion of cellularization (**Fig. 7**). In Arabidopsis, FIS2-PRC2 can regulate a group of type I MADS-box genes in early endosperm development (Köhler *et al.*, 2003; Figueiredo *et al.*, 2015; Zhang *et al.*, 2018). Seeds with dysfunctional MADS genes, such as *AGAMOUS-LIKE 62* (*AGL62*) and *AGL80*, may be aborted due to cellularization defects (Portereiko *et al.*, 2006; Kang *et al.*, 2008). Type I MADS-box genes probably act as negative regulators of cellularization (Pires, 2014), which is opposite to the role that FIE2-PRC2 plays. We previously found that the type I MADS-box genes in rice were associated with syncytium development; their expression being substantially suppressed during cellularization (Chen *et al.*, 2016a). In the present study, we found that expression of many MADS-box genes were disrupted in *osfie2*, whereas *osfie1* displayed limited impacts on these genes (**Fig. 9C and Supplementary Fig. 7**). The results suggested that OsFIE2-PRC2 may suppress the expression of these MADS-box genes, in order to promote cellularization in rice.

Plants that overexpressed *OsFIE1* showed more H3K27me3 marks on the chromatin (**Fig. 2B**), indicating that, as well as OsFIE2 (Nallamilli *et al.*, 2013), OsFIE1 exhibits methyltransferase activity *in vivo*. However, several lines of evidence revealed in the current study suggested that *OsFIE1* is probably not as essential for rice development as is *OsFIE2*. Firstly, in an evolutionary scenario, *OsFIE2* had undergone more intensive purifying selection than *OsFIE1*, as indicated by the lower *dN*/*dS* of *OsFIE2* in comparison with *OsFIE1* (**Fig. 1B**). Secondly, a mutation in *OsFIE1* had only mild effects on seed development. Cellularization in *osfie1* was not affected, whereas it was significantly delayed in *osfie2* (**Fig. 7**). Thirdly, transcriptome analysis revealed that, for the common DEGs identified from *osfie1, osfie2* and the double mutant, additive effects were observed for the genes’ expression (**Fig. 9D-F**). The *osfie1,2* double mutant usually showed the greatest effects, whereas *osfie2* showed only a moderate interference. Expression of these DEGs were affected to a lesser extent when compared with *osfie2* or *osfie1,2*. Finally, yet importantly, we could readily obtain homozygous mutants of *osfie1* from the callus-regenerated plants at the T_0_ generation (**Fig. 6B**), but *osfie2* homozygotes were not available by the same approach, probably due to its lethality. The results suggested that *OsFIE2* are indispensable for regeneration and vegetative growth of rice. Previous studies had found that *OsFIE2* RNAi plants showed phenotypes such as short stature and low seed setting (Nallamilli *et al.*, 2013; Liu *et al.*, 2016), which are similar to the phenotypes exhibited by*OsFIE1*-overexpressors (Zhang *et al.*, 2012; Folsom *et al.*, 2014). We therefore assumed that OsFIE1 and OsFIE2 might compete with each other for members of PRC2. Overexpression of *OsFIE1* may increase the malfunctional OsFIE1-PRC2 but decrease the functional FIE2-PRC2 in plants.

Autonomous endosperm has been observed in FIS2-PRC2 mutants (Ohad *et al.*, 1996; Chaudhury *et al.*, 1997; Kiyosue *et al.*, 1999). However, inconsistent findings have been reported, regarding whether rice PRC2 mutants are able to induce autonomous endosperm (Nallamilli *et al.*, 2013; Li *et al.*, 2014). Our findings suggested that *osfie2* may cause autonomous endosperm, but at a very low frequency (**Fig. 8**). About 2.6% of the emasculated *OsFIE2*^+−^ produced autonomous endosperm. This rate was slightly higher in *osfie1*/*OsFIE2*^+−^ (∼4.9%). If the frequency of autonomous endosperm was all contributed by the *osfie2* haploid (representing half of the female gametes produced by a heterozygote), the frequency of autonomous endosperm by *osfie2* homozygotes would be no more than 10% in rice. This is much lower than that observed in *AtFIE* of Arabidopsis. Nearly 50 percent of female gametes produced by a *FIE*/*fie* heterozygote had multinucleate central cells at six days after anthers removed (Ohad *et al.*, 1996), but in a genetic background dependent manner (Roszak & Kohler, 2011). The findings suggested that, in addition to PRC2 complex, suppression of proliferation of the central cell before fertilization requires other regulators in rice. Recent findings suggested that auxin is a signal molecule common to both Arabidopsis and rice for the induction of autonomous endosperm and early seed development (Zhao *et al.*, 2013; Figueiredo *et al.*, 2015, 2016; Batista *et al.*, 2019). In Arabidopsis, the activation of auxin biosynthesis is negatively regulated by FIS2-PRC2 in the central cell (Figueiredo & Köhler, 2018). Whether rice employs a similar way to block mitotic division of the central cell before fertilization needs to be further elucidated. The evidence provided in the present study suggested that PRC2 defects caused by OsFIE1 and OsFIE2 may not induce activation of auxin biosynthesis in rice before fertilization, as reflected in the low frequency of autonomous endosperm observed in *osfie1* and *osfie2* (**Fig. 8A**).

## Materials and methods

### Plant materials and growth conditions

The rice (Kitaake, *O. sativa* ssp. *japonica*) plants were grown in paddy fields in Yangzhou, Jiangsu Province, China, during the summer season, and in paddy fields in Lingshui, Hainan Province, during the winter season. The plants were managed with addition of standard amounts of water and nutrients, according to local farming practices. The seeds within the spikelets were labeled with marker pens with respect to the date of anthesis. Different-aged caryopses, along with other tissues, were collected for subsequent experiments.

### Phylogenetic analysis and calculation of *dN*/*dS*

The FIE homologs of monocots were identified through BlastP searching, using OsFIE1 and OsFIE2 protein sequences as the queries to search against the protein database of EnsemblPlants (http://plants.ensembl.org/index.html). The homologs obtained were aligned by Clustal Omega (https://www.ebi.ac.uk/Tools/msa/clustalo/). The aligned sequences were submitted to MEGA7 for phylogenetic analysis using the maximum likelihood method. The Ka/Ks program of the TBtools package (Chen *et al.*, 2018b) was used to calculate *dN*/*dS. L. perrieri* was used as an out-group. Coding sequences were used for the calculations.

### Vector constructions and transformation

A gene SOEing (Splicing by Overlap Extention) approach (Horton *et al.*, 2013) was applied to amplify *ChiFIE1, ChiFIEb* and *pOsFIE2::OsFIE1*. The coding sequence of *OsFIE1* and the inter-changed *OsFIEs* were cloned into pENTR/D-TOPO entry vector (Invitrogen) and then recombined into the destination vector, pANIC6A, for the *Ubi*::*OsFIE1* and *Chi-OsFIE*s constructs. The reactions were carried out following the instructions of the manufacturer of the LR Clonase Kit (Invitrogen). The coding sequence of *OsFIE1* was cloned into pCAMBIA1300-GT1 to obtain the *OsGT1*::*OsFIE1* construct using the ClonExpress^®^ II One Step Cloning Kit (Vazyme). The promoter region of *OsFIE2* and the coding sequence of *OsFIE1* were simultaneously cloned into pCAMBIA1300 for the *pOsFIE2*::*OsFIE1* construct using the ClonExpress^®^ II Multi step Cloning Kit (Vazyme). The targets for CRISPR/Cas9 constructs were cloned into BGK032 using a CRISPR/Cas9 vector constructing kit (Biogle). The primers used for vector constructions are listed in Supplementary Table 3. The constructs were then transformed into the embryo-induced calli of rice. The methodologies were as described in our previous report (Chen *et al.*, 2016a).

### DNA extraction and bisulfite sequencing

Genomic DNA was isolated from leaves and endosperm of wild-type rice using the EasyPure Plant Genomic DNA Kit (Transgen). Bisulfite sequencing was performed as described in our previous report (Chen *et al.*, 2016a). Briefly, 1 μg of genomic DNA was treated with sodium bisulfite using an EZ DNA Methylation Kit (Zymo). The treated DNA was used as the templates to amplify the *OsFIE1* and *OsFIE2* segments (150–300 bp; primers are listed in Supplementary Table 3). We then cloned the PCR products into a pEAZY-T3 cloning vector (Transgen). For each amplicon, at least 10 clones were sequenced. The sequencing data were submitted to Kismeth for bisulfite analysis (Gruntman *et al.*, 2008).

### RNA extraction and real-time PCR assay

Total RNA was extracted from different tissues of rice ‘Kitaake’ using Plant RNA Kit (Omega). First-strand complementary DNA was synthesized by HiScript Q-RT SuperMix for qPCR (Vazyme) using the oligo-dT primer. Quantitative real-time qPCR was performed according to the manufacturer’s instructions of the AceQ qPCR SYBR Green Master Mix (Vazyme), using the CFX Connect Real-Time PCR Detection System (BioRad). Three biological replicates were set up for each experiment. The primers used in the experiment are listed in Supplementary Table 3.

### RNA-Seq and differential expression analysis

RNA isolated from caryopses of the WT, *osfie1, osfie2* and *osfie1,2* at 5 DAF was used for RNA-Seq. We set up three biological replicates for WT and *osfie1*, and two biological replicates for *osfie2* and *osfie1,2*. The samples were submitted to Novogene Co. Ltd (Tianjin) for library preparation and sequencing. Briefly, 1.5 μg high-quality RNA per sample was used for library preparation using NEBNext^®^ Ultra™ RNA Library Prep Kit, and index codes were added to each sample. Library quality was assessed on the Agilent Bioanalyzer 2100 system. The clustering of the index-coded samples was performed on a cBot Cluster Generation System using the TruSeq PE Cluster Kit v3-cBot-HS (Illumina). The libraries were sequenced using the Illumina Nova 6000 platform.

The raw fastq reads were processed through in-house perl scripts to remove adapters and low-quality reads. The clean reads were imported into the CLC Genomics Workbench 12.0 and mapped to the reference genome using global alignment mode with the default mapping parameters. The expression values, including total counts, unique counts, TPM and RPKM, were calculated for the identification of DEGs, using the threshold fold-change>2 and FDR < 0.05.

### Nuclear protein extraction and western blotting

The leaves from WT and *Ubi:OsFIE1* were finely ground into a powder with liquid nitrogen. The nuclear protein extraction buffer [20 mM Tris-HCl (pH 8.0), 300 mM NaCl, 0.4 M sucrose, 10 mM MgCl_2_, 1 mM dithiothreitol, 1 mM phenylmethylsulfonyl fluoride, and 5 mM 2-mercaptoethanol] was added with the ratio leaves (g)/buffer (ml) of 1:20 for histone protein extraction. The extracted mixture was sequentially precipitated by sulfuric acid solution (0.25 M), trichloroacetic acid (TCA) (90% 0.25 M sulfuric acid, 10% TCA), and pre-cooled acetone. After removing the supernatant and drying the pellet, an appropriate amount of phosphate-buffered saline (pH 7.0)PBS was added to re-suspend the pellet.

The nuclear proteins were used for western blot analysis. Antibodies used in this study were anti-H3K27me3 (Abcam) and anti-H3 (Abcam). Two independent experiments were performed with two biological replicates for each sample.

### Sectioning and microscopy

The caryopses were fixed and infiltrated in FAA solution, and stored at 4 °C for use. Samples were dehydrated through a graded ethanol series and infiltrated with xylene, and then embedded in resin, sectioned at 2.5 μm, and stained with 0.1% Toluidine Blue. The embryo sac and embryogenesis observation were performed by a previously described method with modification (Hara *et al.*, 2015). Briefly, the caryopses were fixed in FAA overnight. After replacing the solution with 70% ethanol, the samples were incubated overnight, subjected to an ethanol series and washed twice with PBS for 30 min. After treatment with RNase A (100 μg/ml) for 20 h, the samples were stained with DAPI (Sigma, 5 μg/ml) at 4 °C overnight in dark. The samples were then washed with PBS three times each for 3 h, dehydrated by an ethanol series, and washed twice with absolute ethanol for 30 min. After dehydration, the samples were treated with a methanol-methyl salicylate series (2:1, 1:1, and 1:2 (v/v) each for 1 h), washed three times with methyl salicylate for 1 h, kept at 4 °C overnight in methyl salicylate and observed by a laser confocal microscope (Zeiss, LSM710).

### Germination assay

To test the germination of near-mature seeds, the seeds (∼25-30 DAF) produced by the main stem were collected. The panicles were put into a horizontally placed germination bag in growth chamber (14-h-day and 10-h-night settings, 25 °C/20°C day/night temperature). The germination rate was calculated every day. To test the germination ability of different aged seeds, spikelets were marked at the day anthesis, different aged seeds were harvested and put into a germination bag in a growth chamber. The germination rate was investigated at 7 days after imbibition.

### Assessment of the frequency of autonomous endosperm

The spikelets were emasculated one day before flowering. A pollination bag was used to cover the emasculated panicle in case cross-pollination. If a caryopsis was enlarged without pollination at 25 days after the anthesis removed, we regard it as an autonomous endosperm.

## Acknowledgment

This work was supported by National Key R&D Program of China (2016YFD0100902), National Natural Science Foundation of China (31771744, 31571623 and 31701392), Science Fund for Distinguished Young Scholars of Jiangsu Province (BK20180047), and the Priority Academic Development of Jiangsu Higher Education Institutions. We thank Dr. Dongping Zhang for critical reading of the manuscript.

## Figure legends

**Figure 1. Evolutionary analysis of FIE homologs of rice.**

(A). Phylogenetic tree of the FIE homologs of monocots. EnsemblPlants IDs of the homologs were provided. The initials of the accession numbers indicate the origin of the gene. All the rice homolog IDs start with “O”. LPRER, AET, HORVU, BRADI, SORBI, Zm, SETIT, GSMUA and Dr indicates *Leersia perrieri, Aegilops tauschii, Hordeum vulgare, Brachypodium distachyon, Sorghum bicolor, Zea mays, Setaria italica, Musa acuminata* and *Dioscorea rotundata*, respectively. The maximum likelihood method was used for the tree construction.

(B). Violin plot of *dN*/*dS* of different *FIE1* and *FIE2* homologs in the genus *Oryza. L. perrieri* was used as an out-group for the *dN*/*dS* calculations.

**Figure 2. Deleterious effects of ectopically expressed *OsFIE1* on vegetative growth are dosage dependent**.

(A). Phenotype of the wild type (WT) and a representative *OsFIE1*-overexpression line (*Ubi::OsFIE1*) at the heading stage.

(B). Relative expression of *OsFIE1* in leaves of *Ubi::OsFIE1* plants. (C). H3K27me3 was elevated in *Ubi::OsFIE1*.

(D). Phenotype of WT and a *proOsFIE2::OsFIE1* line at the heading stage.

**Figure 3. Ectopic expression of the chimeric *OsFIE* did not result in vegetative defects**

(A). Scheme of the chimeric OsFIEs, achieved by swapping the extra N-terminal tail of OsFIE1 to OsFIE2. The green and yellow bars indicated OsFIE1 and OsFIE2 proteins, respectively.

(B). Confirmation of ectopic expression of the chimeric *OsFIEa* (*Chi-OsFIEa*) and *Chi-OsFIEb* in transgenic plants. 1-qRT and 2-qRT indicated the segments used to distinguish the *OsFIE1*- and *OsFIE2*-origin transcripts. The corresponding positions of 1-qRT1 and 2-qRT are indicated in (A). Three biological replicates were used; the error bars indicated standard deviations.

(C, D). Phenotypes of *Chi-OsFIEa-1* (C) line and *Chi-OsFIEb-1* (D) plants at the heading stage.

(E, F). Tiller numbers (E) and plant height (F) of different transgenic lines of *Ubi::OsFIE1, Chi-OsFIEa* and *Chi-OsFIEb*. Twenty plants for each line were measured; error bars indicate standard deviations.

**Figure 4. Overexpression of *OsFIE1* resulted in reduced seed size**.

(A). Phenotypes of seeds from *Ubi::OsFIE1* and *GT::OsFIE1*.

(B). Confirmation of the expression up-regulation of *OsFIE1*, but not *OsFIE2*, in the caryopses (6 days after fertilization (DAF)) of different *GT1::OsFIE1* transgenic lines.

(C). Phenotypes of the wild-type (WT) and *GT::OsFIE1* plants at the heading stage.

(D-G). Length (D), width (E), thickness (F) and 1000-grain weight (G) of the brown seeds produced by the *GT1::OsFIE1* lines.

**Figure 5. Phenotyping of the *osfie1* mutants.**

(A). Illustration of the targets (1-T1 and 1-T2) that were used for generation of CRISPR/Cas9 mutants.

(B). Phenotype of a wild type (WT) plant and two *osfie1* mutants at the heading stage.

(C). Seed phenotype of WT and *osfie1* mutants. Images from top to bottom are WT, *osfie1-1, osfie1-2* and *osfie1-3*, respectively.

(D-G). 1000-grain weight (D), length (E), width (F), and thickness (G) of the brown seeds produced by the *osfie1* lines.

(H). Reduced dormancy of the *osfie1* mutants.

(I). Dynamic curves of germination of WT, *osfie1-1* and *osfie1-2.*

(J). Germination rate of different-aged seeds of WT, *osfie1-1* and *osfie1-2.*

**Figure 6. Generation of *osfie2* mutant.**

(A). Schematic drawing of the three targets (2-T1, 2-T2 and 2-T3) that were used for generation of CRISPR/Cas9 mutants of *osfie2*.

(B). Cumulative percentage of the not-edited (wild type, WT), monoallele-edited (Hetero) and diallele-edited homozygous (Homo) individuals that were regenerated from *Agrobacterium*-mediated transformation at T_0_ generation. The number of individuals of each genotype are indicated on the bars.

(C). Sequencing of 16 F_2_ individuals derived from *OsFIE2*^+−^.

(D). Cumulative percentage of well-filled, unfilled and unfertilized seeds produced by WT (n = 985), *osfie1-1* (n = 1768), *osfie1-2* (n = 1115), *OsFIE2*^+−^ (n = 4772) and *osfie1*/*OsFIE2*^+−^ (n = 3626).

(H). Caryopses collected from a single panicle of an *OsFIE2*^+−^ plant at 15 days after fertilization (DAF). Two classes of caryopsis were observed: Class 1 that produced starchy endosperm, and Class 2 that produced watery endosperm.

**Figure 7. Early endosperm development of wild type (WT), *osfie1, osfie2* and *osfie1,2*.**

(A-L). Sections of 2 days after fertilization (DAF) (A-D), 3 DAF (E-H) and 4 DAF (I-L) endosperm of WT (A, E, I), *osfie1-1* (B, F, J), *osfie2* (C, G, K) and *osfie1,2* (D, H, L).

(M, N). Sections of 7 DAF endosperm of WT (M) and *osfie2* (N).

(O, P). I_2_-KI staining of 7 DAF endosperm of WT (O) and *osfie2* (P).

(Q-X). Expression of some key genes involved in starch biosynthesis. Three biological replicates were used; error bars indicated standard deviations.

**Figure 8. Autonomous fertilization of *OsFIE2***^**+−**^ **and osfie1/*OsFIE2*** ^**+−**^.

(A). Cumulative percentages of the non-fertilized and autonomously fertilized seeds in wild type (WT), *osfie1-1, osfie1-2, OsFIE2*^+−^ and *osfie1*/*OsFIE2*^+−^. The number of each type of seed was indicated on the bars.

(B, C). Morphology of dried autonomous seeds of *OsFIE2*^+−^ (B) and *osfie1*/*OsFIE2*^+−^ (C).

**Figure 9. Transcriptome analysis of caryopses 5 days after fertilization (5 DAF) of wild type WT, *osfie1, osfie2* and *osfie1,2*.**

(A, B). Venn diagrams of the upregulated (A) and down-regulated (B) genes identified from *osfie1, osfie2* and *osfie1,2* in comparison to WT.

(C). MapMan pathway enrichment analysis of the differentially expressed genes (DEGs). Circle size and colors indicate the log scale of the enrichment.

(D). Heatmap of the expression of DEGs common to *osfie1, osfie2* and *osfie1,2*. Gene expression was indicated by log2(FPKM).

(E, F). Additive effects of *OsFIE1* and *OsFIE2* on expression of storage protein biosynthesis-related genes (E) and photosynthesis-related genes (F).

**Figure 10. Functional divergence between *OsFIE1* and *OsFIE2* with respect to seed development of rice**.

Both OsFIE1 and OsFIE2 are able to interact with other members to form functional OsFIE1- and OsFIE2-PRC2. Whether the same or distinct components are recruited by OsFIE1 and OsFIE2 to form a polycomb complex is not determined. By modulating the H3K27me3 of its target genes, OsFIE2 acts as a positive regulator of endosperm cellularization and may also function in terms of starch filling of seeds. *OsFIE1* is less active on the regulation of cellularization, seed filling and maturation; but is essential for dormancy. It is not clear whether *OsFIE2* functions on dormancy as well. Dashed lines indicated undetermined components or regulation, and line thickness indicated importance for regulation.

**Supplementary Table 1. Differentially expressed genes identified from *osfie1, osfie2* and *osfie1,2*.**

**Supplemenatry Table 2. Additive effects of *OsFIE1* and *OsFIE2* on storage preotein and photosynthesis related genes’ expression.**

**Supplementary Table 3. Primes used in the study.**

---

**Supplementary Figure 1.**
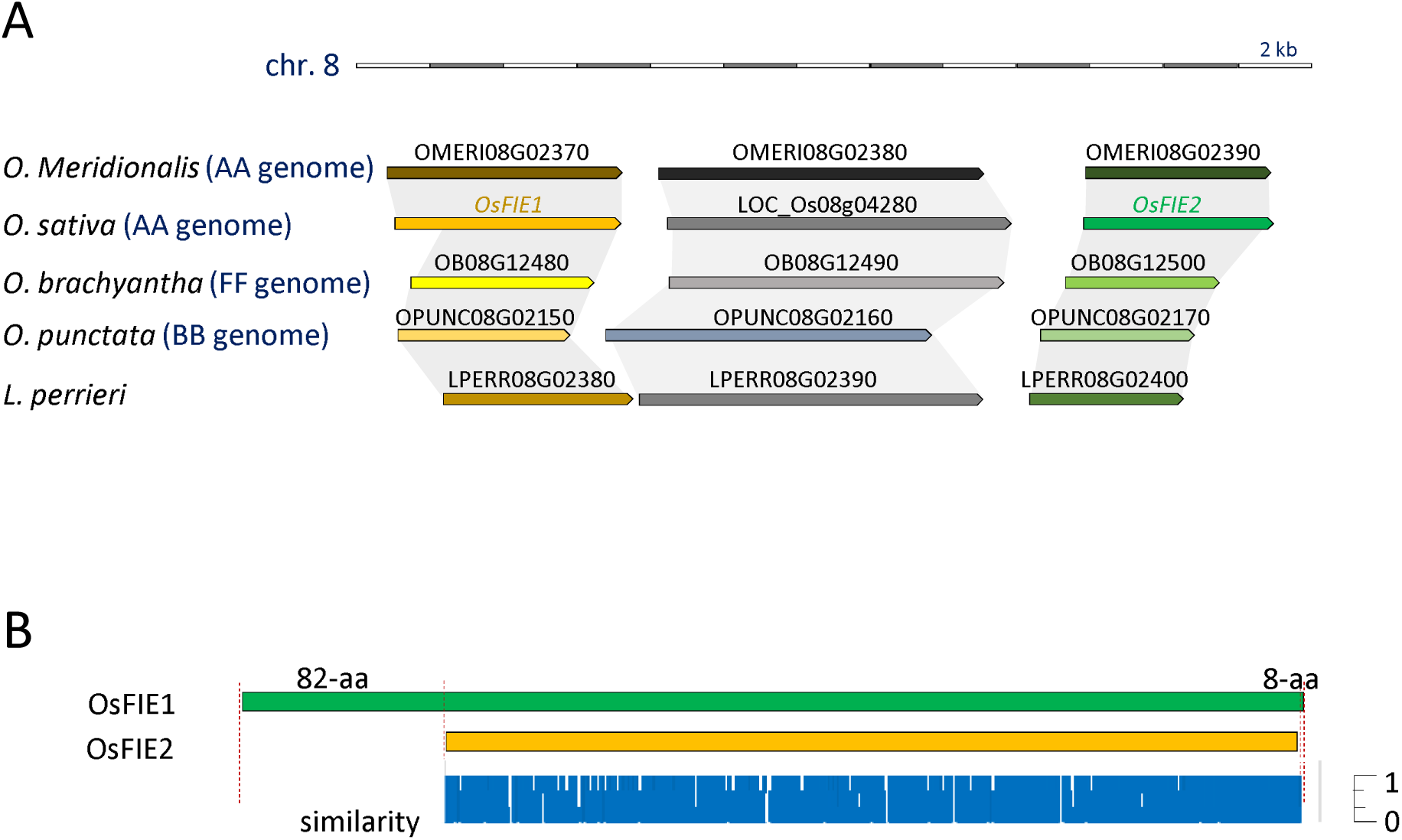
Synteny of the chromosomal segments that containing *FIE* homologs among different Oryzeae species and alignment of OsFIE1 and OsFIE2. (A). Synteny among *O. sativa*, O. *Meridionalis, O. brachyantha, O. punctata* and *L. perrieri* of the chromosomal segment that embedded the duplicated *FIE* homologs. (B). Alignment of OsFIE1 and OsFIE2. OsFIE1 has an extra N-terminal segment that consists 82 amino acid residuals (aa) and an extra C-terminal segment that consists 8 aa. The similarities between OsFIE1 and OsFIE2 at corresponding positions are indicated by blue bars.

**Supplementary Figure 2.**
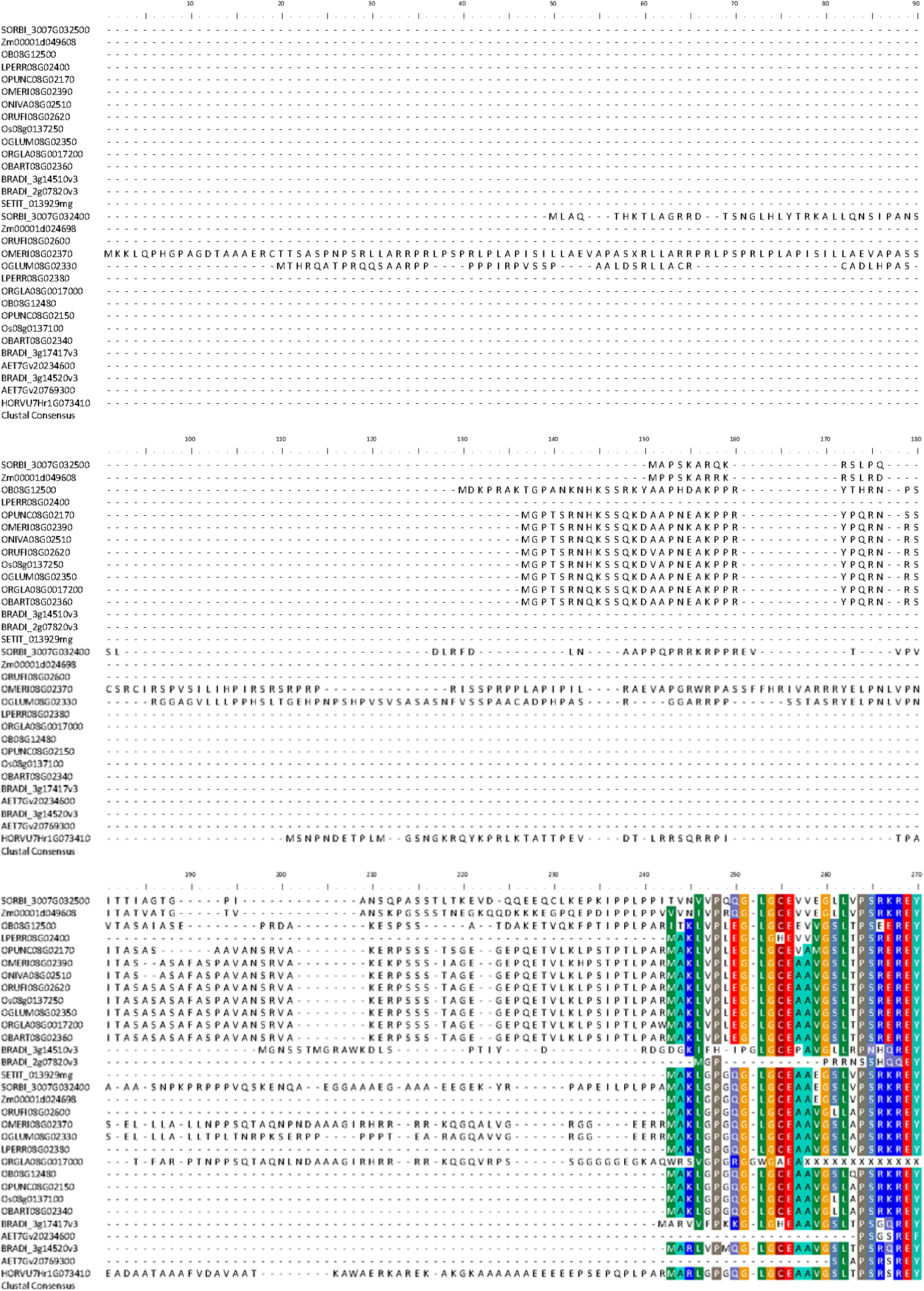
Alignment of the N-terminus of OsFIE1 and OsFIE2 homologs.

**Supplementary Figure 3.**
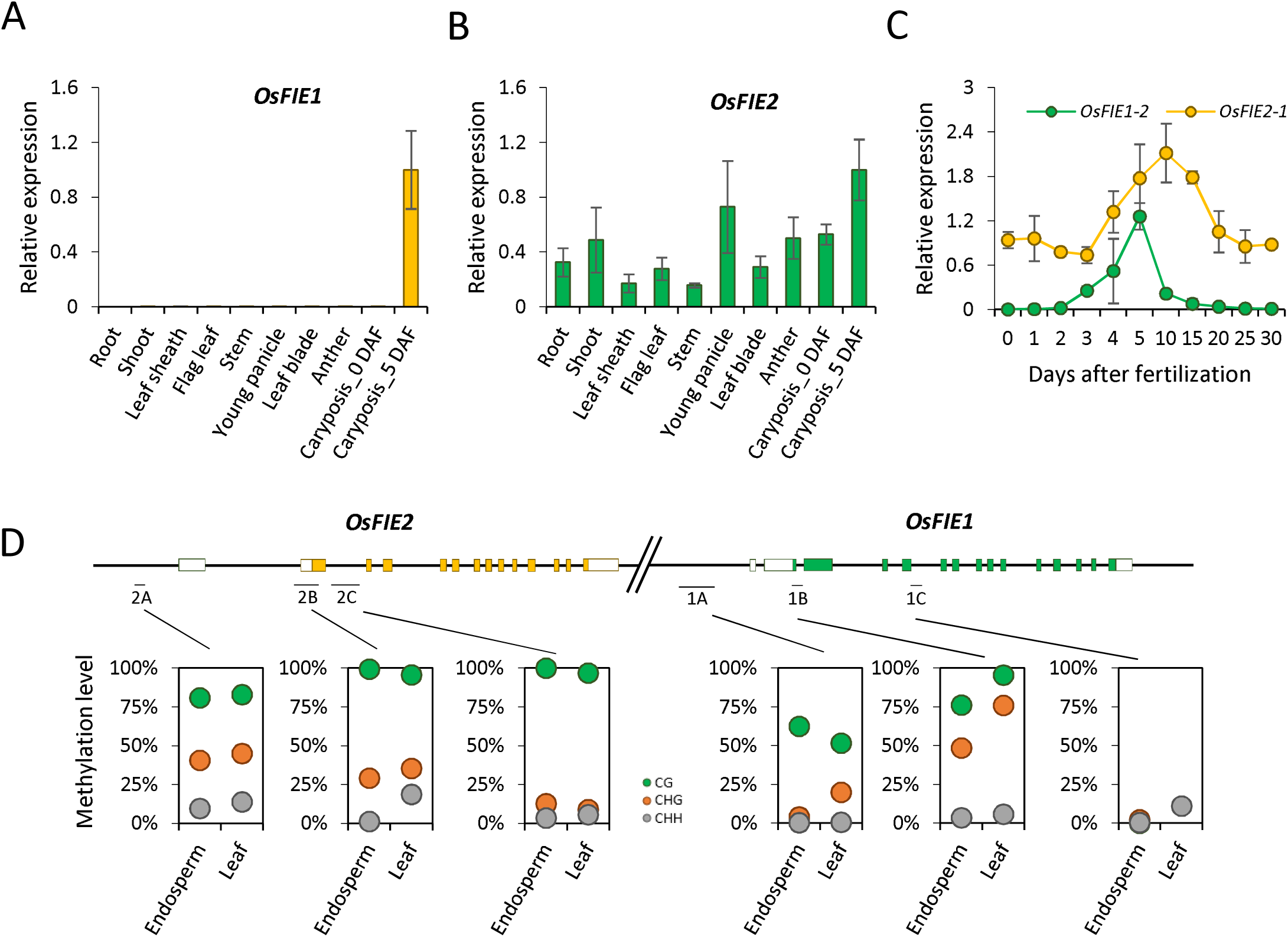
Expression profiles and DNA methylation landscapes of *OsFIE1* and *OsFIE2*. (A, B). Relative expression of *OsFIE1* (A) and *OsFIE2* (B) in different tissues of rice. (C). Relative expression of OsFIE1 and OsFIE2 in different aged caryopsis of rice. (D). DNA methylation level of different segments of *OsFIE1* and *OsFIE2* that indicated on the schematic figure. At least 10 clones of each BS-PCR product were sequenced to estimate the methylation.

**Supplementary Figure 4.**
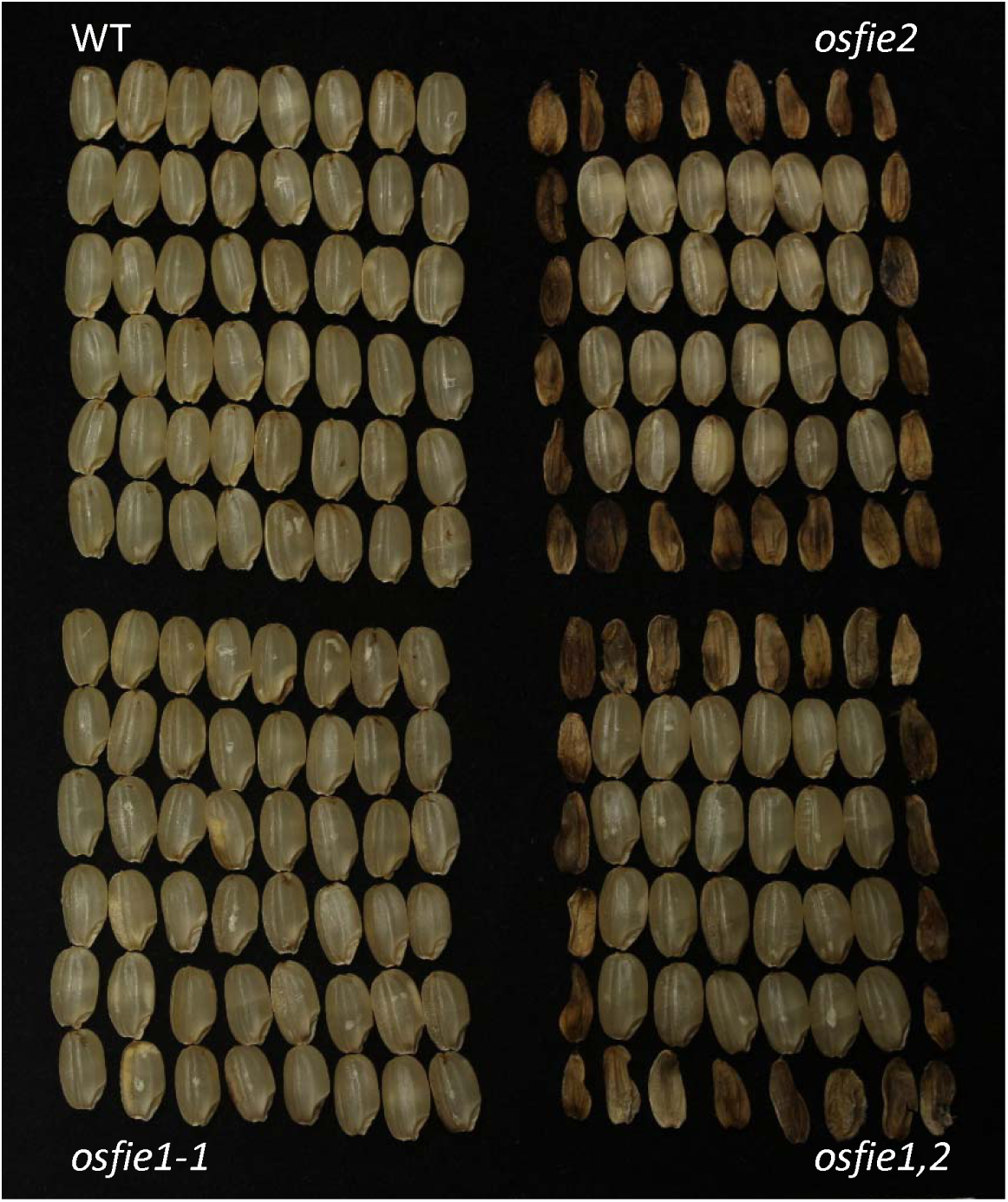
Seed phenotype of WT, *osfie1, OsFIE2*^+−^and *osfie1*/*OsFIE2*^+−^ at mature stages. Class 1 seeds produced by *OsFIE2*^+−^ and *osfie1*/*OsFIE2*^+−^ (central) were resemble to those produced by WT and *osfie1*. Class 2 seeds (peripheral) were enlarged but empty.

**Supplementary Figure 5.**
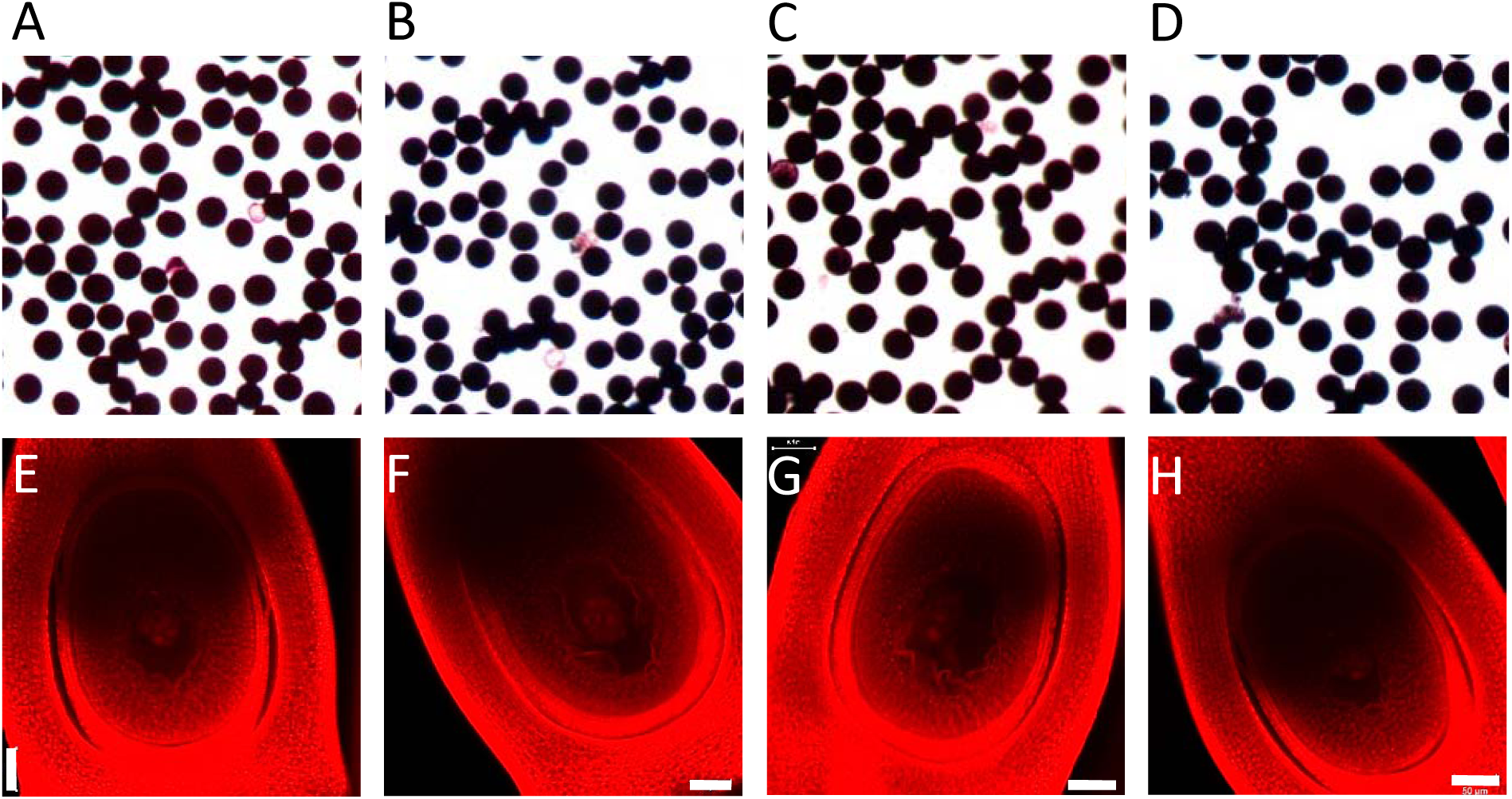
The viability of pollen grains and embryo sac of WT, *osfie1, osfie2* and *osfie1,2*. (A-D). I2-KI staining of the pollen grains of WT (A), *osfie1* (B), *OsFIE2*^+−^ (C) and *osfie1*/*OsFIE2*^+−^ (D). There were no viability differences showed among the pollens of different genotypes. (E-H). CMSL observation of the embryo sacs of WT (E), *osfie1* (F), *OsFIE2*^+−^ (G) and *osfie1*/*OsFIE2*^+−^ (H). A representative embryo sac for each genotype was presented. The embryo sacs generally showed no difference between different genotypes. Bar=50 μm.

**Supplementary Figure 6.**
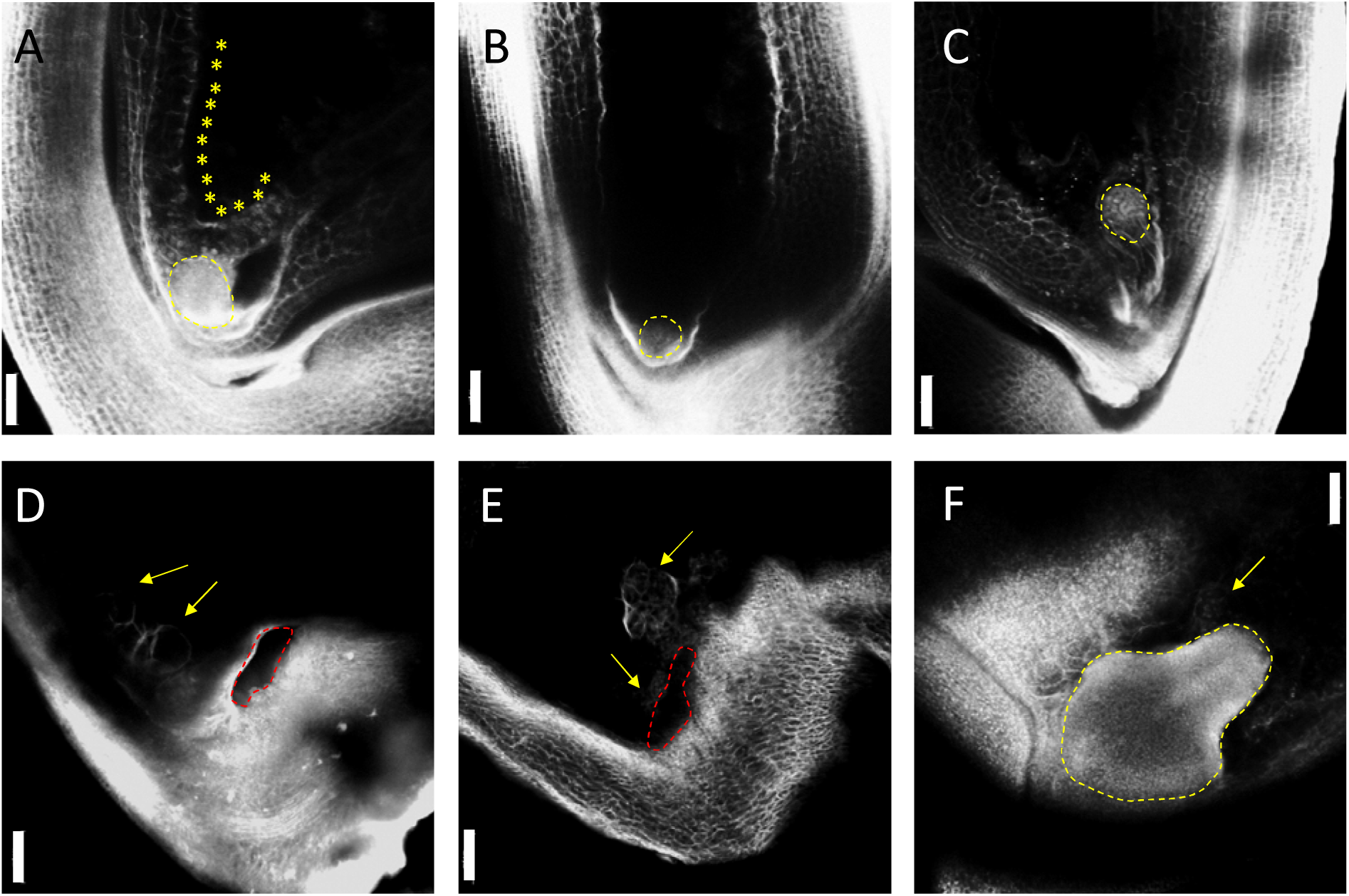
CLSM observation of the embryo development of WT, *osfie2* and *osfie1,2*. (A-C). Embryo development of WT (A), *osfie2* (B) and *osfie1,2* (C) at 4 DAF. Globular embryos are circled by yellow dash lines. The cellularized cells of WT are indicated by stars. Bar=50 μm. (D-F). Observation of the embryos of *osfie2* (D) and *osfie1,2* (E and F) at 15 DAF. The cavities left by degenerated embryos are indicated by red dash lines in (D and E); an arrested yet degenerated embryo of *osfie1,2* is highlighted by yellow dash line. The abnormal endosperm cells are indicated by arrows. Bar=100 μm.

**Supplementary Figure 7.**
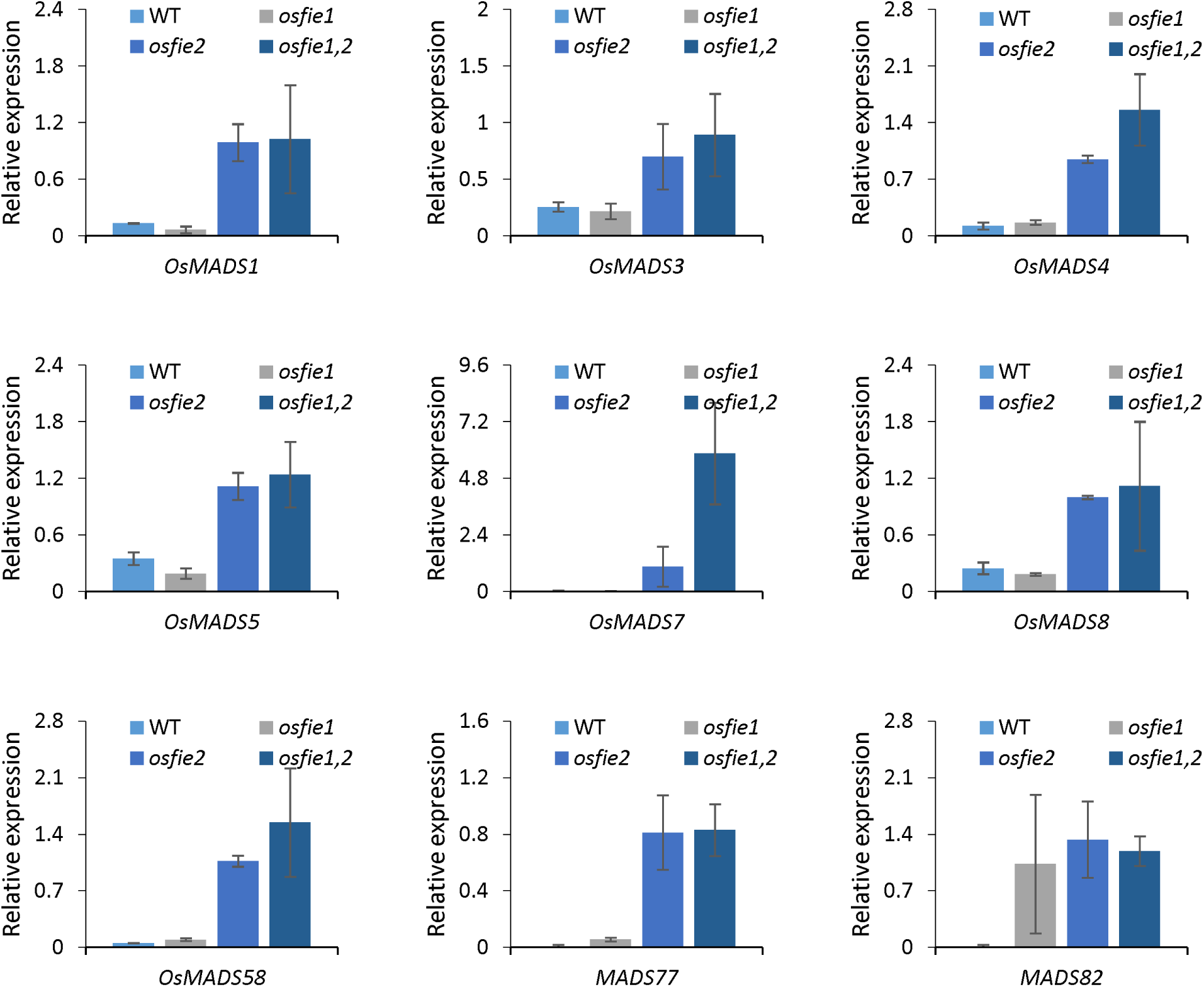
Relative expression of some MADS-box genes in the endosperm of WT, *osfie1, osfie2 and osfie1,2*.

